# Keeping FIT: Iron-mediated post-transcriptional regulation in *Toxoplasma gondii*

**DOI:** 10.1101/2023.11.08.565792

**Authors:** Megan A. Sloan, Adam Scott, Clare R. Harding

## Abstract

Iron is required to support almost all life; however, levels must be carefully regulated to maintain homeostasis. Although the obligate parasite *Toxoplasma gondii* requires iron, how it responds when iron becomes limiting has not been investigated. Here, we show that iron depletion triggers significant transcriptional changes in the parasite, including in pathways that require iron. Interestingly, we find that a subset of *T. gondii* transcripts contain stem-loop structures which have been associated with post-transcriptional iron-mediated regulation in other cellular systems. We validate one of these (found in the 3’ UTR of TGME49_261720) using a reporter cell line. We show that the presence of the stem-loop containing UTR is sufficient to confer accumulation at the transcript and protein levels under low iron. We show that this response is dose and time-dependent and is specific for iron. Using immunoprecipitation, we show that the metabolic enzyme aconitase is capable of binding to mRNA, including *fit*, and that the presence of the *fit* UTR leads to stabilisation of the transcript under low iron conditions. These results demonstrate the existence of iron-mediated post-transcriptional regulation in *Toxoplasma* for the first time, and show that the metabolic enzyme aconitase may have an additional role as an RNA-binding protein.

## Introduction

*Toxoplasma* is a highly successful obligate intracellular parasite with a broad host range. Infection in humans is typically asymptomatic, although in pregnant or immunocompromised people infection can have severe consequences. During infection, *T. gondii* invades and replicates within multiple cell types–exposing the parasite to rapidly changing levels of the key nutrients required to support replication. One of these essential nutrients is iron, *Toxoplasma gondii* requires iron to power cellular metabolism and replication (Bergmann *et al*., 2020; Pamukcu *et al*., 2021a). We have found that *Toxoplasma* can detoxify excess iron by storing it within a membrane-bound compartment through the action of the vacuolar iron transporter (VIT), and that this storage is required to protect cells from excess iron (Sloan, Aghabi and Harding, 2021; Aghabi *et al*., 2023) However, the parasite is likely to be exposed to limiting iron levels during infection as some tissues have low levels of available iron, and mammalian hosts restrict iron availability during infection (Reinert *et al*., 2019; Morel *et al*., 2022). This means the parasite must sense and respond to changing levels of iron to maintain pathogenesis, however the mechanisms underlying this remain unknown.

Due to the importance of iron across kingdoms of life, regulation of iron uptake, usage and storage is highly conserved, (Wang and Pantopoulos, 2011; Martínez-Pastor and Puig, 2020; Gao and Dubos, 2021; Fontenot and Ding, 2023). However, mechanisms used to respond to iron stress are frequently distinct between organisms and range from control at the transcriptional (Fontenot and Ding, 2023), post-transcriptional and protein (Shin *et al*., 2013) levels. These pathways allow cells to regulate key proteins in response to low iron to promote survival.

One of the best studied mechanisms for iron-mediated regulation in mammalian cells is the post-transcriptional IRE/IRP system. Transcripts of genes involved in cellular iron homeostasis (such as the transferrin receptor, TfR, the main pathway of cellular iron uptake) contain short stem-loop regions (SLRs), called iron-response elements (IREs)(Campillos *et al*., 2010; Sanchez *et al*., 2011). In low iron conditions, the metabolic enzyme aconitase (also called iron-regulatory protein 1, IRP1) loses its iron-sulphur (FeS) cluster cofactor. This abrogates its enzyme activity, and instead aconitase moonlights as a RNA-binding protein where it binds to the IREs present in the untranslated regions (UTRs) of specific genes (Hentze *et al*., 1987; D. M. Koeller *et al*., 1989; Philpott, Klausner and Rouault, 1994). Crucially, the location of the IRE within the transcript determines whether aconitase binding results in repression or promotion of translation. Aconitase binding in the 3’ UTR of genes acts to stabilise transcripts and promotes protein production (D M Koeller *et al*., 1989; Erlitzki, Long and Theil, 2002). Whereas binding in the 5’ UTR, found in transcript of the iron storage protein ferritin, is thought to block access of the translation apparatus (Hentze *et al*., 1987; Sanchez *et al*., 2007; Garza *et al*., 2020), inhibiting translation. This system allows cells to respond to iron stress by upregulating iron import (via TfR) and downregulating storage, leading to greater iron availability for cellular processes.

The aconitase/IRE system has been reported to regulate iron response in several organisms beyond mammals, including invertebrates and bacteria (Alén and Sonenshein, 1999; Tang and Guest, 1999; Lind *et al*., 2006). In plants, the role of aconitase in iron-response remains unclear; in *Arabidopsis thaliana* aconitases can bind RNA (Marondedze *et al*., 2016), however this did not appear to directly modulate ferritin abundance (Arnaud *et al*., 2007), possibly explained by functional redundancy between the three aconitases. Iron-mediated regulation beyond model organisms is less well understood. In the kinetoplast *Trypanosoma brucei,* an RNA binding protein binds and stabilises the transcript of the parasite transferrin receptor under low iron to promote iron uptake (Carbajo *et al*., 2021). Similar systems have been proposed in the protozoan parasites *Trichomonas vaginalis* (Calla-Choque *et al*., 2014; León-Sicairos *et al*., 2023) and *Giardia duodenalis* (Plata-Guzmán *et al*., 2023). Within the apicomplexan family, there is some evidence of a functional IRE system. In *Plasmodium*, the causative agent of malaria, IRE-like structures have been identified and shown to interact with parasite aconitase (Loyevsky *et al*., 2001, 2003). However, these experiments were performed *in vitro* using bacterially purified proteins, and the existence of a functional system in cells was not established.

Here we perform RNAseq on iron-deprived parasites and find global transcriptional changes, including alterations in iron-dependent pathways. To identify possible mechanisms for parasites responses to low iron, we identify *in silico* IRE-like sequences in the *T. gondii* transcriptome. Selecting a predicted IRE from the 3’ UTR of a predicted, but unstudied iron transporter, TGME49_261720, which we name here FIT. We show that this IRE-like sequence is required and sufficient to confer iron-responsivity to a reporter gene. Interestingly, we further show that this IRE-structure binds to parasite aconitase, and that the transcript is stabilised under low iron conditions. This is the first example of a functional IRE-like sequence in *T. gondii,* and provides evidence for a role of post-translational gene expression regulation in *T. gondii* iron homeostasis.

## Results

### The transcriptional response to iron deprivation

Transcriptional responses to iron deprivation are common across organisms (Andrews, Robinson and Rodríguez-Quiñones, 2003; Li *et al*., 2008; Gao and Dubos, 2021). To determine how iron deprivation impacts the transcriptome of *T. gondii,* we deprived parasites of iron by infecting primary human cells (HFF), pretreated for 24 h with the iron chelator Deferoxamine (DFO). After 24 h, parasites were mechanically released, filtered to remove host debris and bulk RNA-sequencing performed (**Figure 1A**). We observed 1197 transcripts with higher abundance in parasites cultured in low iron conditions compared to untreated parasites, whilst just 44 were downregulated (log_2_fold change £-2, adjusted *p*-value (padj) < 0.05) (**Table S1**). This bias towards upregulation of transcripts was unexpected and was not seen upon oxidative stress induction (Augusto, 2021) or upon downregulation of an essential metabolic regulator (Li *et al*., 2023).

**Figure 1.**
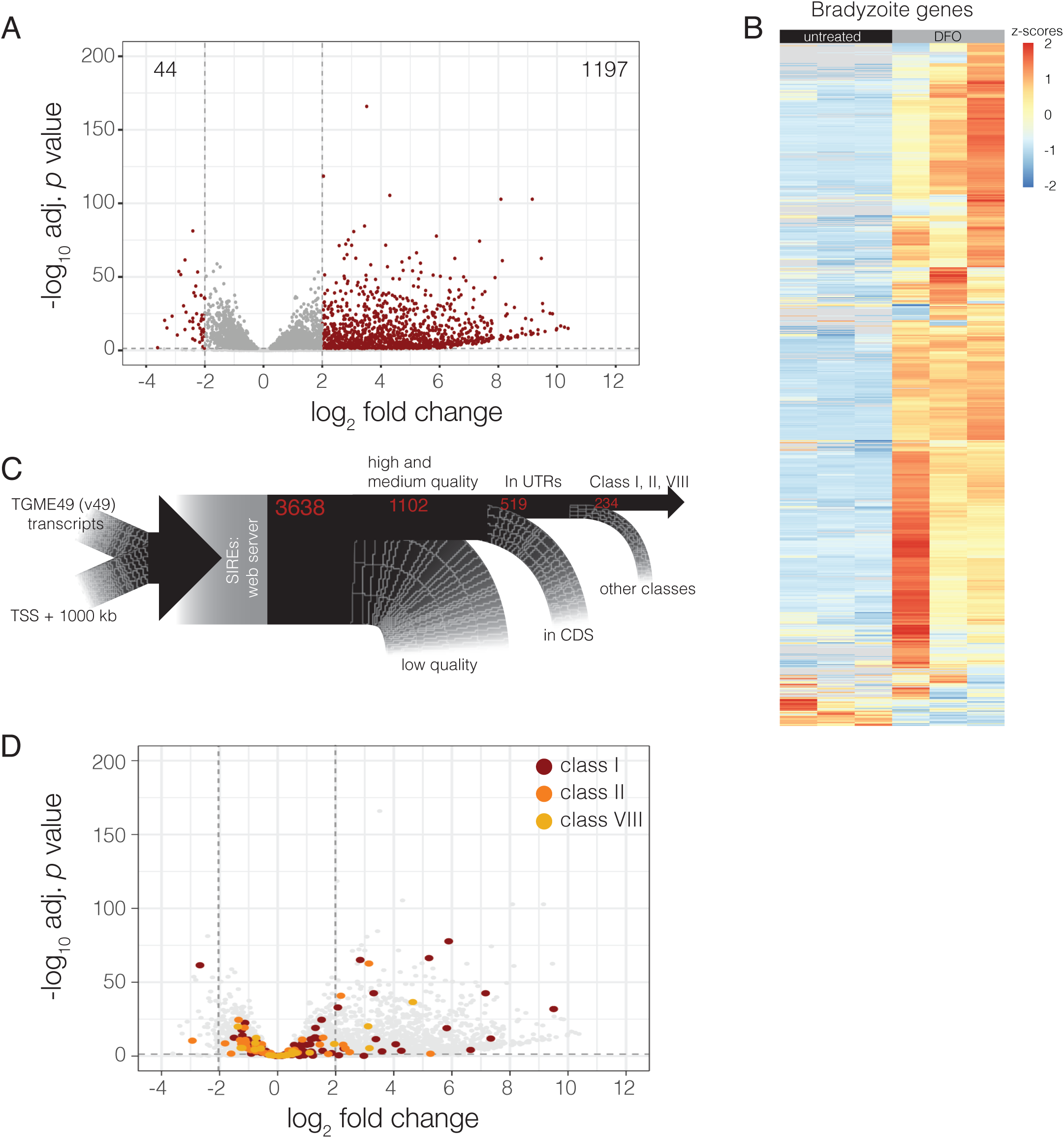
The *T. gondii* transcriptional response to iron deprivation. **A.** Volcano plot from RNAseq data comparing RHΔKu80 cultured in 100 µM DFO for 24 hours to standard conditions. Adjusted p-values from the Wald test with Benjamini and Hochberg correction. Cut-offs shown with dashed lines are p-adj < 0.05 and log_2_ fold change of >2 or < -2. **B.** Heatmap from RNAseq dataset depicting z-scores of genes shown to be upregulated in bradyzoite forms of *T. gondii* by Waldman *et al*. 2020 (adjusted p-value < 0.05 and log_2_ fold change > 2), in DFO treated and untreated parasites. **C.** Schematic showing the filtering process to identity transcripts containing IRE-like sequences in the *T. gondii* transcriptome. **D.** Volcano plot as in (**A)** where transcripts with IRE-like sequences in classes I (red), II (orange) and VIII (gold) are highlighted. See text for more details.

We next examined if iron deprivation impacted transcript abundance of genes whose protein products are predicted to contain iron sulphur (FeS) clusters, using the *in silico* prediction tool MetalPredator (Valasatava *et al*., 2016). Of 60 FeS containing genes, we found 27 were significantly differentially expressed in our dataset (padj < 0.05) with 67% of these transcripts being down regulated (**Table S2**), representing a significant (p < 0.001, Fisher’s exact test) enrichment of predicted FeS-containing genes in the downregulated genes. *T. gondii* encodes three FeS biosynthesis pathways (Pamukcu *et al*., 2021b), of these, the impact of iron deprivation appears minimal on the cytosolic (CIA) and apicoplast (SUF) FeS synthesis pathways with 1/8 and 2/9 genes modestly regulated (log_2_fold changes <1, padj < 0.05, **Table S3**). The largest affect was seen in the mitochondrial FeS synthesis pathway, with 8/12 genes in the ISC pathway differentially regulated including downregulation of the FeS scaffold ISCU1 (TGME49_237560) and the proposed FeS carrier NFU4 (TGME49_212930). The FeS co-chaperone MGE1 (TGME49_265220) and ferredoxin MFdx1 (TGME49_236000) also down regulated, though fold changes were modest. However, ISD11 and NFS1, which provide sulphur to clusters, were upregulated. Supporting this, many of the downregulated FeS-containing genes are predicted to be mitochondrially localised (Barylyuk *et al*., 2020) (**Table S2**).

Of the upregulated transcripts, we saw bradyzoite stage specific genes including: BAG1 (log_2_ fold change 3.2, p-adj = 1.94E-10), ENO1 (log_2_ fold change 5.1, p-adj = 6.3E-18) and LDH2 (log_2_ fold change 6.4, p-adj = 6.87E-29), supporting a recent observation (Zhu *et al*., 2023), . We also observe that the majority of transcripts found to be upregulated in bradyzoites in previous work (Waldman *et al*., 2020) were also upregulated in iron-deprived parasites (**Figure 1B, Table S3**). This suggests that iron deprivation, similar to other nutrient deficiencies (Bohne and Roos, 1997; Fox, Gigley and Bzik, 2004; Ihara and Nishikawa, 2014), can promote *Toxoplasma* differentiation.

Given the strong, apparently loosely targeted effect of iron deprivation on the *T. gondii* transcriptome, we decided to search for potential downstream post-transcriptional regulation which could allow the parasite to respond specifically to low iron.

### The *T. gondii* transcriptome contains IRE-like sequences

We identified transcripts containing putative iron response elements in the *T. gondii* using the online RNA structural prediction tool SIREs (Campillos *et al*., 2010) (**Figure 1C**). The algorithm identified 3638 sequences of interest (**Table S4**), based on homology to known IREs. Many were assigned a ‘Low’ quality score and were not considered further, leaving a total of 1102 genes with IRE-like sequences (**Table S5**). Most functionally validated IRE elements in other systems are present in the untranslated regions of transcripts (Sanchez *et al*., 2011). There were 343 genes with predicted IRE-like sequences in the 5’ UTR and 176 genes with IRE-like sequences in the 3’ UTR (**Table S6**). IREs can be assigned to 18 classes based on sequence conservation to experimentally validated sequences shown to bind aconitase *in vitro*, however the majority of validated IREs are usually in classes I, II or VIII (Campillos *et al*., 2010). As such we focused on the 234 genes with UTR-localised IRE-like sequences from these motif classes (I – 116, II - 92, VIII – 26, **Table S7**).

We then asked how many of the transcripts containing IRE-like sequences were differentially regulated in our RNA-seq dataset. We found that only 142 (60%) of IRE-like sequence containing genes were regulated by iron availability (p-adj < 0.05, **Table S7**, **Figure 1D**). As such, our list of putative IRE-like sequences likely contains false positives.

In order to test whether the IRE-like sequences can impact gene expression we chose to focus on a class VIII IRE-like sequence (5’-gctgcctccgtgtcagggtagacgagagaaa-3’) (**Figure S1A**) in the 3’ UTR of a previously unstudied gene TGME49_261720 which was conserved between T. gondii genomes (**Figure S1A**). TGME49_261720 encodes an essential ZIP-domain containing protein (Sidik *et al*., 2016) and is a putative plasma-membrane localised zinc/iron transporter (Barylyuk *et al*., 2020). For these reasons, we renamed the gene FIT.

### The *fit* 3’ UTR is sufficient to render a reporter gene iron responsive

To determine whether the predicted IRE in the *fit* transcript is sufficient to confer iron-responsivity, we cloned *fit* 3’ UTR immediately downstream of the fluorescent protein tdTomato under a constitutive promoter (tubulin-tdTomato-fit). We also constructed a control line with tdTomato upstream of the *dhfr* 3’ UTR (tub-tdTomato-dhfr) which does not contain an IRE-like sequence and we did nt expect to be iron responsive. We integrated these cassettes into the *uprt* locus of previously developed RH *T. gondii* parasites constitutively expressing mNeonGreen fluorescence protein in the *ku80* locus from the *sag1* promotor with a *sag1* 3’ UTR (Aghabi *et al*., 2023) and confirmed by PCR (**Figure S1B-E**). Interestingly, replacement of the *dhfr* UTR with *fit* resulted in significantly lower tdTomato signal in untreated parasites (**Figure S1F**), demonstrating the influence of the 3’ UTR on protein expression, as previously observed (Smith *et al*., 2022).

To determine if the *fit* UTR conferred iron responsivity, we pre-treated host cells for 24 hours with the iron chelator deferoxamine (DFO) before infecting with our reporter and control lines. At 24 hours post infection we collected cells and quantified mNeonGreen and tdTomato fluorescence by flow cytometry (**Figure 2A and B**). DFO treatment did not significantly affect mNeonGreen fluorescence in either line, and, in the tdTomato*_dhfr_* line, iron deprivation had no effect on tdTomato fluorescence. However, DFO treatment led to an approximately 2-3 fold increase in tdTomato fluorescence in the tdTomato*_fit_* line (**Figure 2B**). We quantified the ratio of tdTomato/mNeonGreen fluorescence and normalised to the untreated lines, this showed a significant (*p* < 0.0001, one way ANOVA) increase in the ratio in DFO treated parasites, compared to the tdTomato*_dhfr_* line (**Figure 2C**). Interestingly, there was no significant difference between fluorescence ratio in parasites treated with excess iron, demonstrating that this response is only seen in low iron conditions.

**Figure 2.**
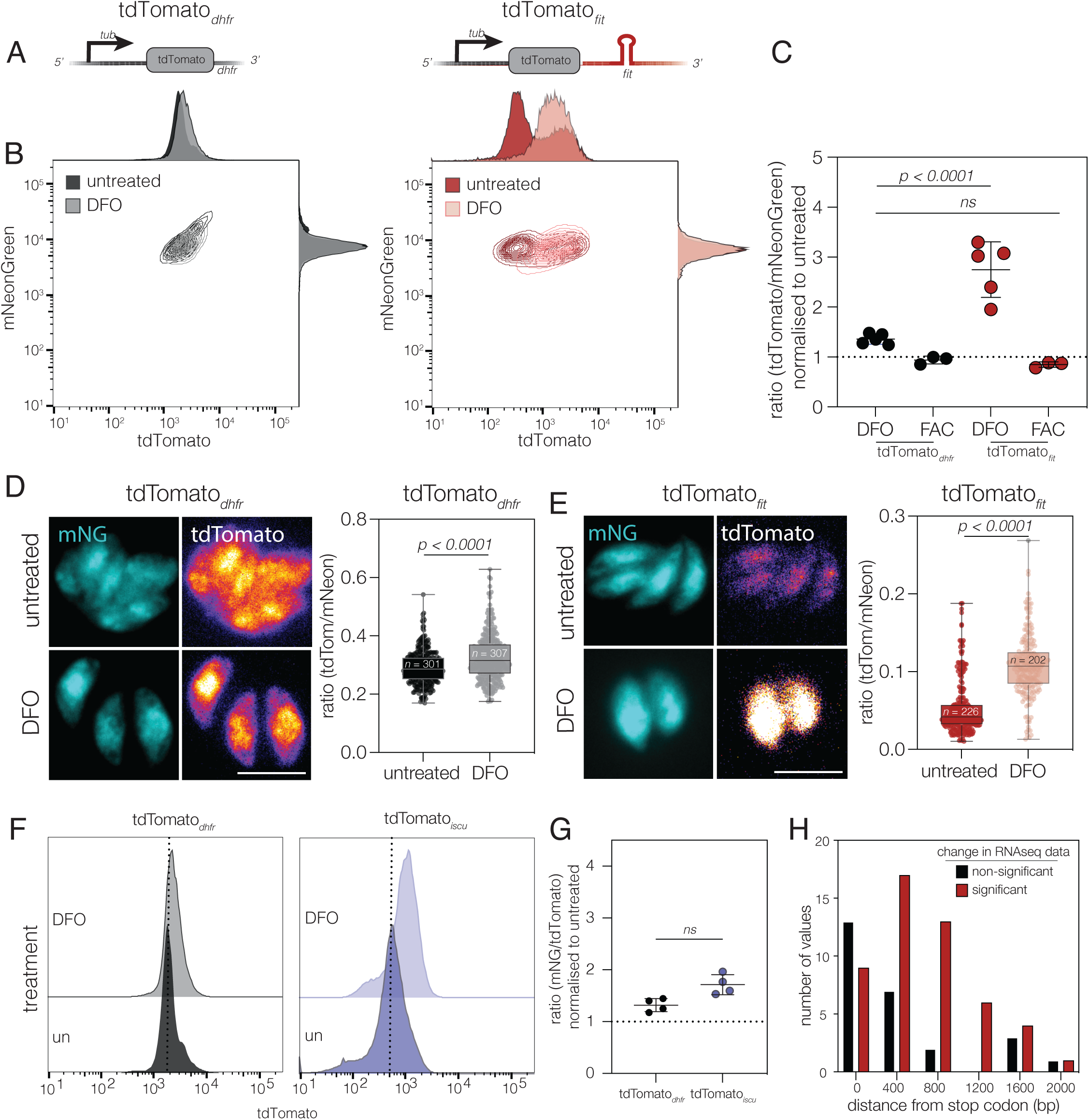
The *fit* 3’ UTR is sufficient to render a reporter gene iron responsive. **A.** Schematic of the tdTomato*_dhfr_* cassette and tdTomato*_fit_* cassette. Both reporters were integrated into the *uprt* locus in a ΔKu80:mNeonGreen parasite line. **B.** Contour plots of mNeonGreen and tdTomato expression in untreated treated with 100µM DFO for 24 hours (measured by flow cytometry) for both reporter lines. Representative of at least 5000 parasites from a single experiment. **C.** Plot showing the mean tdTomato:mNeonGreen fluorescence ratio in tdTomato*_dhfr_* and tdTomato*_fit_* parasites treated with either 100µM DFO or 2.5 mM ferric ammonium citrate (FAC), normalised to untreated parasites. Plot shows data from 5 independent experiments. *p* values compared to tdTomato*_dhfr_*by one way ANOVA with Dunnett’s correction **D.** Immunofluorescence showing mNeonGreen and tdTomato expression in untreated and DFO treated (24 hours, 100 µM DFO) tdTomato*_dhfr_* parasites with a violin/box plot showing the quantification of tdTomato:mNeonGreen fluorescence from vacuoles for from two biological replicates (*n* indicated on graph), *p* values from unpaired t test. Scale bar 5 µm **E.** Immunofluorescence showing mNeonGreen and tdTomato expression in untreated and DFO treated (24 hours, 100µM DFO). Scale bar 5 µm. tdTomato*_fit_* parasites with a violin/box plot showing the quantification of tdTomato:mNeonGreen ratio as above. **F.** Representative overlapping histograms showing tdTomato fluorescence in the tdTomato*_iscu_* reporter line both untreated and after 24 hours of treatment in 100 µM DFO. **G.** Plot showing the mean tdTomato:mNeonGreen fluorescence ratio in untreated tdTomato*_iscu_* and those treated with 100 µM DFO, normalised to untreated parasites. Data shown from 4 independent experiments. *p* value from unpaired t test. **H.** Histogram showing the frequency of differentially regulated transcripts with IRE-like sequences in their 3’ UTRs in RNAseq data. Transcripts were binned (bin size = 400 bp) based on distance of the IRE-like sequence from the stop codon.

To confirm these results, we also assessed tdTomato fluorescence by microscopy and confirmed that there was an increase in the ratio of red:green fluorescence in the tdTomato*_fit_* line upon DFO treatment, but not in the tdTomato*_dhfr_*line (**Figure 2D and 2E**).

Given the impact of the *fit* 3’ UTR on gene expression we chose to test a second 3’ UTR with a predicted IRE-like sequence. For this we chose another gene involved in parasite iron metabolism ISCU (TGME49_237560) the mitochondrial FeS pathway scaffold protein. *iscu* is predicted by SIREs to have a medium quality, class 13, IRE-like sequence in the 3’ UTR (5’- catgctgaatgcatgccatgtaccatgaacat -3’). As above we produced an *iscu* reporter line (tub-tdTomato-iscu) and confirmed genomic integration by PCR (**Figure S1D**) and flow cytometry (**Figure S1E**) where it showed an intermediate level of fluorescence. We then quantified the ratio of tdTomato fluorescence in untreated and DFO treated tdTomato*_iscu_* parasites as above. While we saw a slight increase in tdTomato fluorescence upon DFO treatment (**Figure 2F**) this change was not statistically significant (**Figure 2G**). As such this IRE-like sequence is not likely to be functional. Other than a difference in the nucleotide sequence between the *fit* and *iscu* IRE-like sequences, we also noted that they occupy different positions in the respective 3’ UTRs. The *fit* IRE-like sequence is relatively central in the UTR (position 1016/2144 bp), whereas the *iscu* sequence starts closer to the 3’ end of the UTR (position 1149/1278 bp). To examine whether IRE position may be important for function, and therefore offer a hypothesis as to the difference in responsiveness between the *fit* and *iscu* reporters, we plotted the distance from the stop codon of predicted IRE-like sequences in 3’ UTRs against whether that transcript was differentially abundant in our RNAseq dataset (**Figure 2H**). From this we observed that transcripts with IRE-like sequences between 600-1400 bp from the stop codon were more likely to be differentially abundant in iron deprived parasites compared to the control. Therefore, IRE-like positioning may be important to function, and this may account for the lack of response seen in the tdTomato*_iscu_* line.

### Fit responsiveness is dose dependent and specific to iron chelation

Given the responsiveness of the tdTomato*_fit_* line, we then further validated the reporter line to determine whether this response is dosage dependant. We first repeated our initial experiments by pretreating host cells with concentrations between 0-500 µM of DFO for 24 hours before infection with the tdTomato*_dhfr_* and tdTomato*_fit_* reporter parasites for 24 h, then quantifying tdTomato fluorescence by flow cytometry (**Figure 3A**). As previously observed tdTomato fluorescence in the tdTomato*_dhfr_* was not significantly altered by any tested concentration of DFO, however we observed a dose-dependent increase in tdTomato in tdTomato*_fit_* parasites (**Figure 3B**).

**Figure 3.**
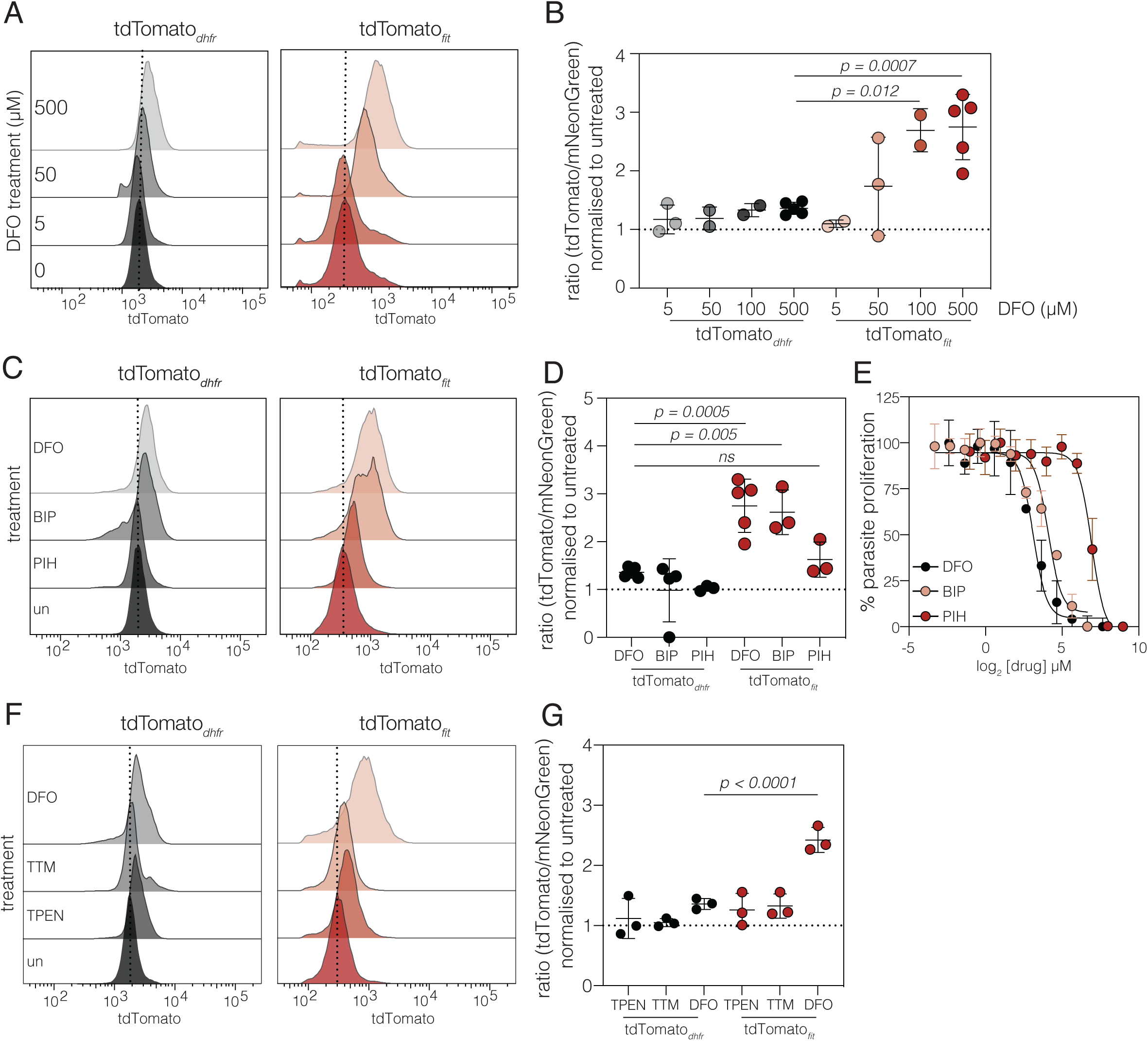
*Fit* responsiveness is dose dependent and specific to iron chelation. **A.** Representative overlapping histograms showing tdTomato fluorescence in the tdTomato*_dhfr_*and tdTomato*_fit_* reporter lines treated with indicated DFO concentration for 24 hours. **B.** Quantification of change of tdTomato:mNeonGreen ratio. *p* values from one way ANOVA with Dunnett’s correction, compared to tdTomato*_dhfr_* + 500 µM DFO **C.** Representative overlapping histograms showing tdTomato fluorescence, measured by flow cytometry, in the tdTomato*_dhfr_* and tdTomato*_fit_* reporter lines treated with 100 µM DFO, 100 µM BIP or 100 µM PIH for 24 hours. **D.** Mean tdTomato:mNeonGreen fluorescence ratio for three independent experiments. *p* values from one way ANOVA with Dunnett’s correction, compared to tdTomato*_dhfr_* + DFO. **E.** Fluorescence growth assay for tdTomato parasites cultured in increasing concentrations of iron chelators. Points show mean of three independent experiments ± SEM. **F**. Representative overlapping histograms showing tdTomato fluorescence, measured by flow cytometry, in the tdTomato*_dhfr_* and tdTomato*_fit_* reporter lines treated with 100 µM DFO, the zinc chelator 5 µM TPEN or Cu chelator 25 µM TTM for 24 hours. **G.** Mean tdTomato:mNeonGreen fluorescence ratio for three independent experiments. *p* values from one way ANOVA with Dunnett’s correction, compared to tdTomato*_dhfr_*+ DFO.

Following this we then asked whether the tdTomato*_fit_*reporter responses are consistent when treated with different iron chelators. To this end we treated both tdTomato*_dhfr_* and tdTomato*_fit_* parasites with two further membrane-permeant iron chelators, the Fe^2+^ chelator, 2,2’-bipyridine (BIP) and the Fe^3+^ chelator pyridoxal isonicotinoyl hydrazone (PIH). Interestingly, while the response to BIP was indistinguishable from DFO (**Figure 3C and D**), there was no change in tdTomato fluorescence upon treatment with PIH. To investigate this further, we quantified the effectiveness of the three chelators in inhibiting parasite growth (**Figure 3E**) and found that DFO showed the greatest inhibition of parasite replication (EC_50_ 8.68 uM (95% C.I. 6.63-11.31 µM)), with BIP also showing high potency (EC_50_ 17.05 µM, 95% C.I. 11.9 – 24.13 µM) while PIH was significantly (*p* = 0.046, Sum of squares F test) less able to prevent parasite replication (EC_50_ 121.01 µM, 95% C.I. 97.6 – 148.3 µM), suggesting that it is less able to remove parasite-available iron than DFO or BIP in these conditions.

Given that several metal iron transporters are able to transport multiple metal species, e.g. the *Plasmodium* ZIPCO likely has roles in the transport of both iron and zinc (Sahu *et al*., 2014), we sought to confirm that the response seen was specifically due to removal of iron, we also tested two other metal chelators, *N,N,N’,N’*-tetrakis-(2-pyridylmethyl)ethylenediamine (TPEN), a broadly active metal ion chelator with high affinity for Zn^2+^ (Schaefer-Ramadan *et al*., 2019) and Tetrathiomolybdate (TTM), a selective copper chelator (Alvarez *et al*., 2010). Treatment with either of these chelators did not lead to an increase in the tdTomato:mNeonGreen ratio (**Figure 3F and G**), suggesting that the response seen is specific to iron chelation and is not a generalised response upon removal of other metals.

### Kinetics of iron response

To determine the kinetics of the response to iron in our reporter line, cells pretreated with DFO were infected for the indicated times before parasites were mechanically removed. To determine the timing of the response at the transcript level, we performed qRT-PCR on reverse transcribed RNA taken from the parasites (**Figure 4A**). We saw no significant change in the tdTomato transcript abundance (relative to actin) in the tdTomato*_dhfr_* cells. However, in the tdTomato*_fit_* line there was an increase in transcripts detectable from 9 hours (*p* = 0.0849, mixed effects test with Tukey’s multiple comparisons test), which then was significantly increased at 18 hours of treatment (*p* = 0.0169). However, by 24 h (the peak of tdTomato protein abundance) mRNA levels had fallen back to untreated levels. This pattern in regulation was similar to that of the endogenous *fit* transcripts (**Figure 4B**), though the magnitude of the increasing transcript abundance was more modest – possibly reflecting the difference in promoter strength. We also examined protein expression over the same time course using flow cytometry (**Figure 4C**). As at the transcript level, there was no significant change in tdTomato fluorescence was observed in the tdTomato*_dhfr_* cells at any timepoint. However, in the tdTomato*_fit_* line, we saw a gradual increase in tdTomato signal, starting at 18 h post infection with a further increase at 24 h. Interestingly, the peak of increased transcript level was seen earlier than the peak of protein expression. These results show that mRNA levels peak prior to protein, as would be expected. It also suggests that this response to iron deprivation relies on new protein synthesis, rather than a reduction in the degradation at the protein level in the cell.

**Figure 4.**
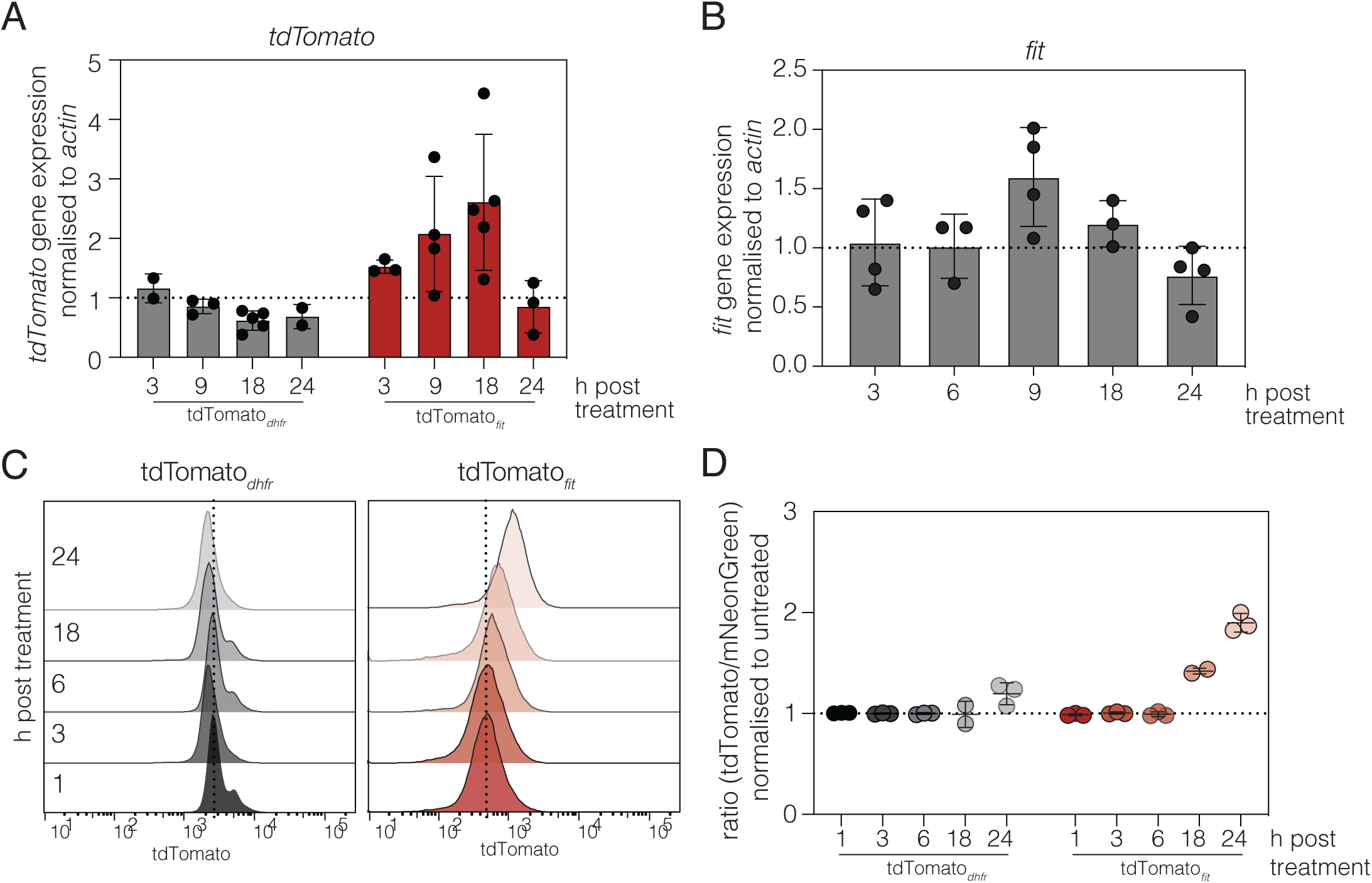
Kinetics of tdTomato*_fit_* iron-responsiveness. qRT-PCR on the relative abundance of *tdTomato* (**A**) and *fit* (**B**) transcripts (normalised to actin) in the tdTomato*_dhfr_* and tdTomato*_fit_*reporter lines after treatment with 100 µM DFO, compared to untreated parasites for indicated time. Points represent a single experiment, bars at meanLJ±LJSD. **C.** Representative overlapping histograms showing tdTomato fluorescence, measured by flow cytometry, in the tdTomato*_dhfr_* and tdTomato*_fit_* reporter lines treated with 100 µM DFO over a 24-hour time course. **D.** Plot showing the mean tdTomato:mNeonGreen fluorescence ratio for the experiments described in (**C).**

### Removal of the IRE-like element from the *fit* 3’ UTR reduces iron responsiveness

As described above, we found a predicted IRE-like element in the *fit* 3’ UTR. To determine if this small region was important for conferring iron-responsivity, we deleted an 18 nucleotide sequence from the predicted IRE (bases 2-25 of the predicted sequence), to create a tdTomato*_fit_*_ΔIRE_ line (**Figure 5A**). Deletion of the IRE has previously been shown to abolish IRE function in other transcripts (Garber and Pudek, 2014; Volz, 2021). Using this line, we assayed transcript abundance in the tdTomato*_fit_*_ΔIRE_ reporter line at 18 hours of treatment, the peak of response from the above time course (**Figure 4A**). The tdTomato*_fit_* line had significantly more tdTomato transcripts (*p* = 0.0019, one way ANOVA with Tukey’s correction) than the tdTomato*_dhfr_* line (**Figure 5B**). However, tdTomato transcript abundance for the tdTomato*_fit_*_Δ*IRE*_ was not significantly different the tdTomato*_dhfr_*line (*p* = 0.44, one-way ANOVA with Tukey’s correction). We also confirmed this at the protein level at 24 h post infection. As shown above, the presence of the *fit* UTR induced a strong increase in tdTomato expression (**Figure 5C**), compared to the control tdTomato*_dhfr_* line (*p* = 0.0023, one-way ANOVA with Dunnett’s multiple comparisons test) (**Figure 5D**), while removal of the IRE-like element (while not affecting total tdTomato fluorescence in untreated cells) prevented the responsiveness upon iron depletion. This suggests that the iron-responsiveness of the tdTomato*_fit_* reporter is dependent on the IRE-like element in the *fit* 3’ UTR.

**Figure 5.**
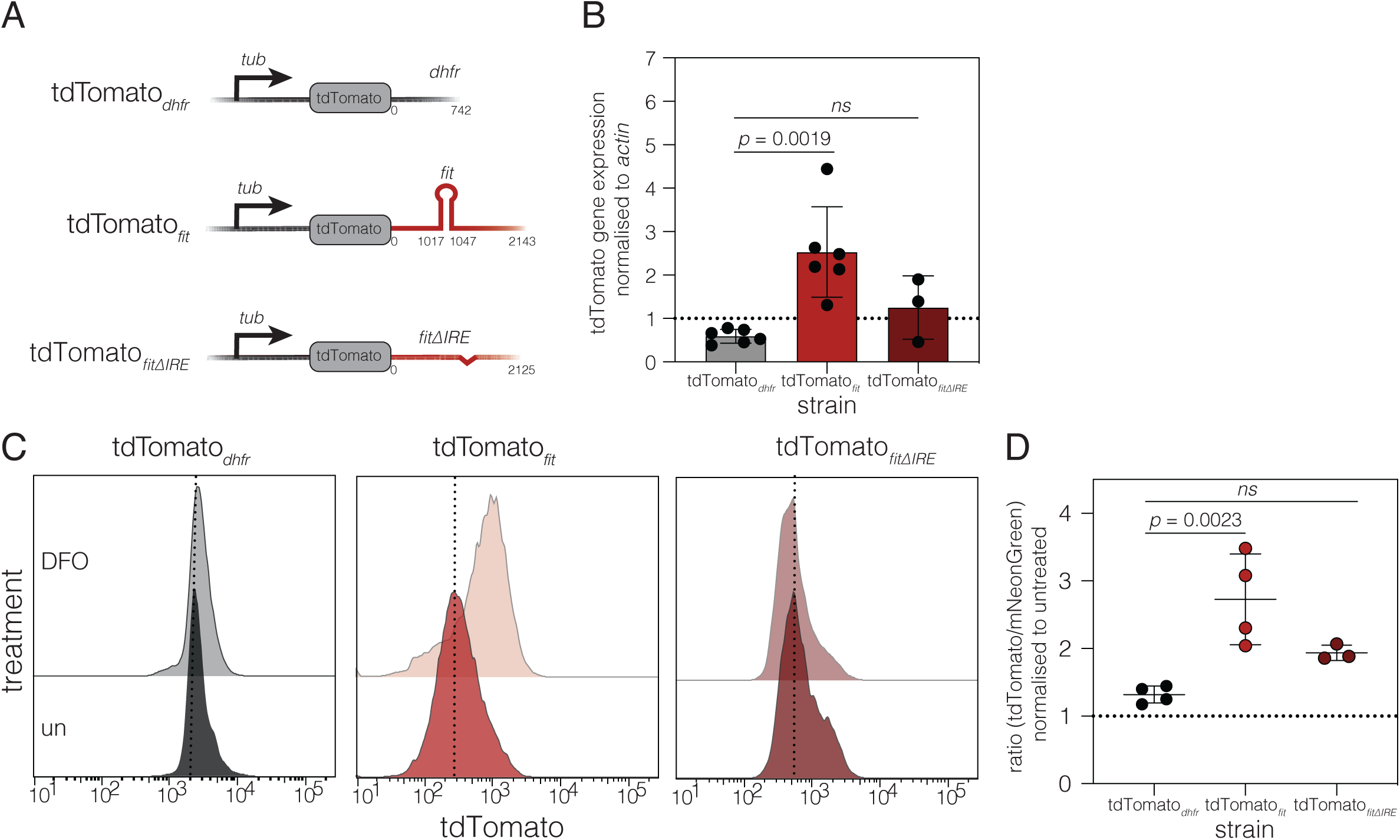
Removal of the IRE-like element from the *fit* 3’ UTR reduces iron responsiveness. **A.** Schematic of the iron reporter cassettes integrated into the *uprt* locus. **B.** qRT-PCR on the relative abundance of *tdTomato* transcripts (normalised to actin) in the the tdTomato*_dhfr_*, tdTomato*_fit_* and tdTomato*_fit_*_ΔIRE_ reporter lines after 18 hours treatment with 100 µM DFO, compared to untreated parasites. Bars at meanLJ±LJSD, *p* values from one way ANOVA with Tukey’s correction. **C.** Representative overlapping histograms showing tdTomato fluorescence in the tdTomato*_dhfr_*, tdTomato*_fit_* and tdTomato*_fit_*_ΔIRE_ reporter lines, treated with 100 µM DFO for 24 hours. **D.** Mean tdTomato:mNeonGreen fluorescence ratio, *p* values from one way ANOVA with Dunnett’s correction, compared to tdTomato*_dhfr_* + DFO.

### Aconitase is associated with *fit* mRNA in low iron conditions

The single aconitase of *T. gondii* (ACN, TGME49_226730) shows high sequence similarity (**Figure S2A**) and predicted structure (**Figure S2B**) with the cytosolic mammalian aconitase, including key residues known to be important for binding the catalytic iron-sulphur cluster and RNA-binding (Walden *et al*., 2006). To investigate the role of aconitase in the response to iron, we made use of a parasite line with aconitase tagged with a Ty epitope (ACN-Ty) (**Figure 6A**). ACN-Ty localised to the mitochondrion, apicoplast and cytosol (**Figure 6B**) as previously described (Pino *et al*., 2007), and localisation was not significantly affected by iron depletion. Parasites expressing mNeonGreen-Ty (Aghabi *et al*., 2023) were included as a control (**Figure 6B**). It has previously been reported that aconitase requires FeS-binding to enable enzymatic activity (Oexle, Gnaiger and Weiss, 1999; Goncalves *et al*., 2008; Gunawardena *et al*., 2016). To determine if iron depletion affected *T. gondii* aconitase enzymatic function, activity was quantified by NADPH accumulation as previously described (Quirós, 2018a). Iron depletion by DFO treatment led to a small but reproducible and significant reduction in enzymatic activity (p< 0.0001, paired t-test, **Figure 6C**), as indicated by a decrease in NADPH in treated parasites. We included a control sample of filtered, uninfected host cells which showed no measurable host cell aconitase is carried over in our parasite samples.

**Figure 6.**
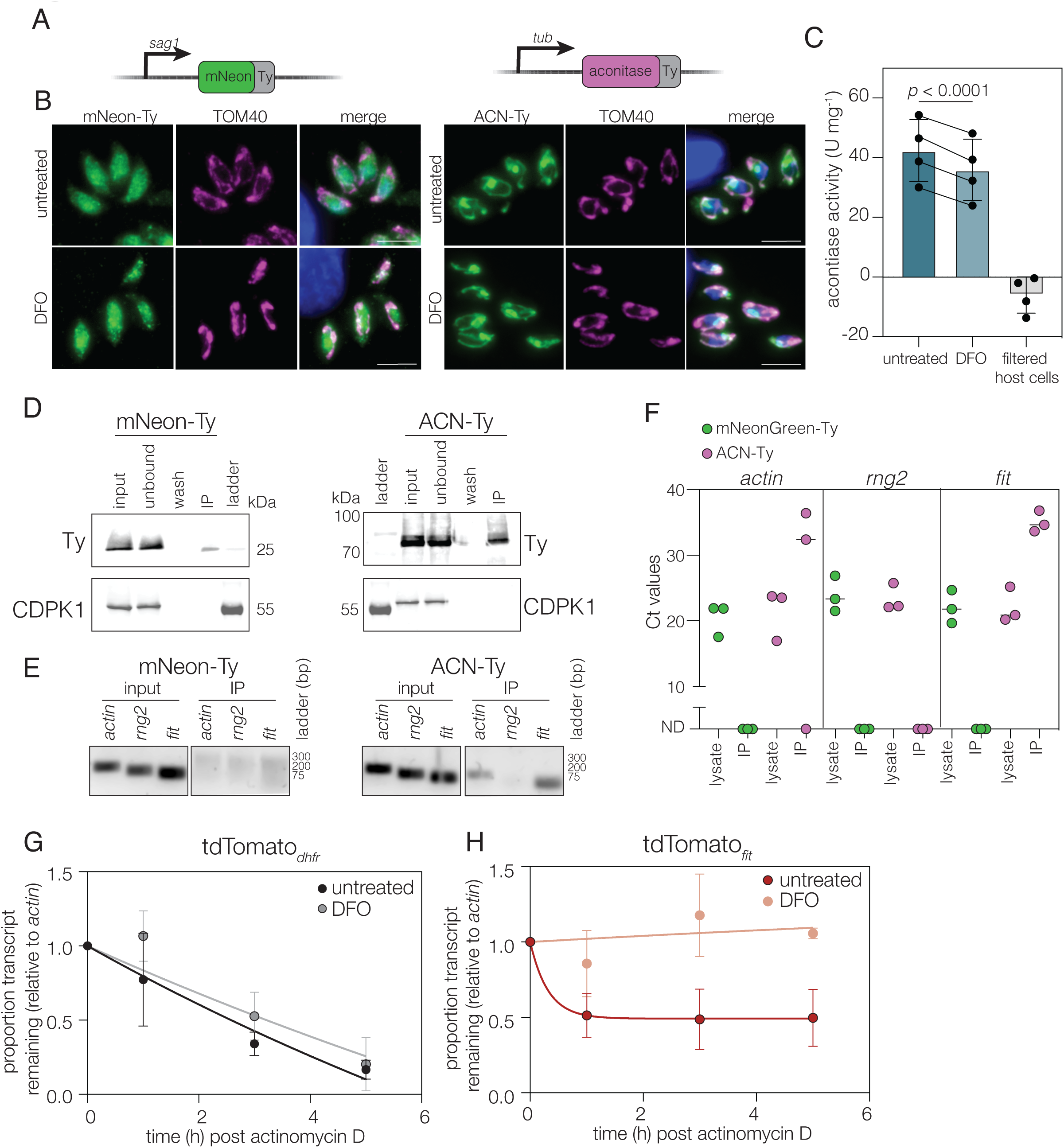
Aconitase binds to the *fit* 3’ UTR. **A.** Schematic of tagging scheme in ΔKu80:mNeonGreen-Ty and ACN-Ty parasites. **B.** Immunofluorescence of ΔKu80:mNeonGreen-Ty and RHΔKu80, ACN-Ty parasites grown in standard culture or 100 µM DFO for 24 hours. Anti-TOM40 included as a mitochondrial marker and DAPI as DNA marker. Scale bars 5µm. **C.** Aconitase activity assay of untreated or DFO treated parasites. Results from 4 independent experiments, ± SD. Lysed and filtered host cells are included to demonstrate minimal carryover of host aconitase activity. *p* value from paired t test. **D.** Blots showing the immunoprecipitation of mNeonGreen-Ty and ACN-Ty from *T. gondii* parasites. CDPK1 included as cytosolic control. Wash – the output of the 6^th^ and final washing step. CDPK included as an experimental control. Representative of three independent replicates **E.** DNA-agarose gel showing qPCR products amplified from reverse transcribed RNA from either starting lysates or RNA co-precipitated with mNeonGreen-Ty and aconitase-Ty proteins. PCR amplicons for *actin* (expected size: 188 bp), *fit* (expected size: 116 bp) and *rng2* (expected size: 146 bp). **F.** qRT-PCR results from Ips performed in (**D**). Points represent average Ct values from individual biological replicates, performed in triplicate, lines at median. ND – not detected. **G.** Untreated or DFO-treated tdTomato*_dhfr,_*treated intracellularly with actinomycin D and qPCR performed. Results normalised to *actin* and the mean of three independent replicates, ± SD. **H.** Untreated or DFO-treated tdTomato*_fit,_* treated intracellularly with actinomycin D and qPCR performed. Results normalised to *actin* and the mean of three independent replicates, ± SD.

Aconitase presence in the cytoplasm would potentially allow the protein to interact with mRNAs. Given this, and the conservation of aconitase RNA-binding role in several clades, we investigated whether *T. gondii* aconitase can bind mRNA. To this end, parasite ACN-Ty was immunoprecipitated from parasites cultured in a low iron environment for 24 hours. As a control, mNeonGreen-Ty (Aghabi *et al*., 2023) was also immunoprecipitated (**Figure 6D, Figure S3B and C**). RNA was then extracted from elutants (and parasite lysates as a loading control) and reverse transcribed to allow identification of associated transcripts by qPCR. Here, we included *actin* as a highly transcribed mRNA and *rng2*, a structural protein of the apical complex (Katris *et al*., 2014), as a control transcript which has a comparable expression level to *fit* within our transcriptomics dataset, but did not show altered expression (**Figure S3C**) and did not contain a predicted IRE. Protein-associated cDNAs were quantified by qRT-PCR (**Table S9**) and visualised on a DNA agarose gel (**Figure 6E**). From ACN-Ty-associated cDNAs we detected transcripts by qRT-PCR from both *actin* and *fit*, but not *rng2* (**Figure 6F, Table S8**) and we were not able to detect any transcripts associated with mNeonGreen-Ty. Together, this suggests that *fit* transcripts are associated with aconitase in *T. gondii* cultured under low iron conditions.

### Iron deprivation increases *fit* transcript stability

Aconitase binding to IRE-sequences in transcripts has been shown to lead to stabilisation of the mRNA, resulting in increased protein production (Owen and Kühn, 1987; Casey *et al*., 1988; Tang and Guest, 1999). Having identified *T. gondii* aconitase as a potential *fit* mRNA-binding protein during iron deprivation, we sought to determine the effect of iron deprivation on *fit* transcript stability. To this end, we measured transcript abundance after blocking new transcription with actinomycin D, as previously described (Heaslip *et al*., 2011). Transcript levels of tdTomato*_dhfr_* fell over the timecourse, in both untreated and low iron conditions, reaching approximately 17% of the starting level after 5 hours of treatment with no significant difference (*p* = 0.19, extra sum of squares F-test) (**Figure 6F**). tdTomato*_fit_*showed a similar pattern in untreated cells, with a more rapid transcript depletion followed by a plateau between 1 and 5 h. However, under low iron conditions we saw a significant (*p* < 0.0001, extra sum of squares F-test) stabilisation of the tdTomato*_fit_* transcript with transcript abundance remaining around 100% even after 5 hours of actinomycin D treatment (**Figure 6G)**. We believe that this stabilisation leads to the increased mRNA abundance we observed, and later to an increase in protein levels (**Figure 4**). Given the binding of aconitase to specific transcripts and the role of the IRE sequence, this work provides evidence for the presence of a post-transcriptional iron-response pathway in *T. gondii*.

## Discussion

Here we show for the first time the existence of iron-mediated post-transcriptional responses in *T. gondii.* The ability of cells to respond to iron availability is conserved across the tree of life, however the mechanistic details of regulation are often species-specific. In *T. gondii*, iron deprivation leads to a substantial upregulation of transcripts, with over 13% of the genome significantly upregulated (log_2_ fold change >2), and few genes downregulated. Along with changes in genes predicted to encode iron-bound proteins and their synthesis pathways, we see evidence that iron deprivation leads to an upregulation of genes associated with the in bradyzoite stage (e.g., BAG1, LDH2 and ENO1) (Waldman *et al*., 2020). Iron deprivation now joins an extensive list of cellular stresses able to trigger stage conversion in *Toxoplasma*. Recent work is now uncovering the upstream molecular pathways involved in mediating stage conversion (Licon *et al*., 2023), however the pathway of how loss of iron directly triggers the stage conversion remains unknown.

The lack of apparent specificity in the transcriptional response led us to believe there were further, post-transcriptional layers of regulation to ensure that *T. gondii* could respond to iron deprivation. Using a sequence-based search algorithm (Campillos *et al*., 2010), we were able to identify genes containing iron response element-like sequences. One limitation of this approach is that these predictions are made based on data from human and mouse cells; *T. gondii* are highly divergent from model opisthokonts (White and Suvorova, 2018) and so while these predictions are likely to be an overestimate, we cannot exclude that functional but divergent IREs are also present. Additionally, the short sequence-based search algorithm utilised cannot consider the full context of transcripts (e.g. folding or secondary structures) and may also return ‘false positive’ sequences. Our data suggests that beyond these limitations, both the IRE class, and its positioning within the UTR, may be important in responsiveness, and future work will seek to more precisely understand which features are required.

The iron response seen here was dynamic, mRNA accumulation was detected between 9-18 h post treatment, while protein levels increased between 18-24 h post treatment. We did not extend treatment beyond 24 h, as between 24-48 h we saw a sharp drop in parasite viability (results not shown). However, mRNA accumulation was transient, with levels returning to below the untreated by 24 h. While this explains the otherwise surprising finding that *fit* is significantly downregulated in our 24 h-treated RNAseq (**Figure S3A**), we do not yet know the mechanisms underlying this repression. We found that removal of 18 nucleotides from the *fit* 3’ UTR mostly abrogated the iron-responsiveness, suggesting that this motif plays a key role. Whilst it is possible that removal of these nucleotides altered the mRNA folding, we confirmed that removal of the predicted IRE did not affect transcript stability or eventual protein abundance under normal growth conditions. Although not statistically significant, the *fit*ΔIRE UTR did show generally greater iron-responsivity than the control, suggesting that other mRNA-structures may contribute, although the IRE appears essential for the magnitude of the response.

We then sought to determine whether, as established in other systems, aconitase can bind the *T. gondii* IRE-like sequences and impact gene expression. *T. gondii* encodes a single aconitase gene required for the TCA cycle (MacRae *et al*., 2012; Harding *et al*., 2020) which conserves all the predicted RNA-binding residues. By immunoprecipitating aconitase from *T. gondii* in low iron conditions, we were reproducibly able to amplify *fit* mRNA, occasionally the highly abundant *actin*, but not the control *rng2* bound to the protein. This provides the first evidence that *T. gondii* aconitase shares a role as an RNA-binding protein, with a potential role in the post-transcriptional response to low iron. We further show that upon iron deprivation, the presence of the *fit* 3’ UTR acts to prevent degradation of the transcript. We believe it is likely this increased stability which leads to the accumulation of transcript, followed by protein levels. In mammalian systems, aconitase binding to IRE-sequences has been shown to stabilise the TfR transcript, leading to protein upregulation (DeRusso *et al*., 1995; Kato *et al*., 2007). Here we cannot yet say definitively that stabilisation of *fit* transcript is due to aconitase binding, however the data we have points in this direction.

One important unanswered question in this work is the role of FIT within the cell. FIT encodes a zinc and iron permase (ZIP) domain containing protein which is predicted by required for growth (Sidik *et al*., 2016) and to localise to the plasma membrane (Barylyuk *et al*., 2020). These features suggest it may be involved in iron uptake by the parasite, however future work will investigate the role and regulation of FIT In summary, we have found that an IRE-like sequence in the 3’ UTR of a proposed iron transporter confers iron responsivity to a reporter gene. This responsivity is specific to iron depletion, and occurs first by stabilising the mRNA transcript, leading to accumulation of the protein. Iron-responsivity requires an 18 nucleotide stretch within the predicted IRE. We show that aconitase is capable of binding to the *fit* transcript, suggesting that this system works in a similar manner to the well-studied mammalian IRE/IRP system.. Future work will seek to define the sequence constraints for this binding and to investigate the biological consequences of such a system in *T. gondii*.

## Methods

### *T. gondii* and host cell maintenance

*Toxoplasma gondii* tachyzoites were grown in human foreskin fibroblasts (HFFs) cultured in Dulbecco’s modified Eagle’s medium (DMEM) maintained at 37°C with 5% CO_2_ that was supplemented with either 3% (D3) or 10% heat-inactivated foetal bovine serum (D10) (FBS), 2 mM L-glutamine and 10 μg ml^-1^ gentamicin.

### Construction of reporter parasite lines

The tdTomato_DHFR_, tdTomato_FIT_ and tdTomato_FITDIRE_ reporter lines were generated by transfection (Donald and Roos, 1995) of 50 mg of plasmid containing Cas9 and sgRNA for the *uprt* locus as previously described (Aghabi *et al*., 2023) along with 100 mL of PCR product containing the relevant reporter gene cassette into the constitutively expressing RHDku80::mNeonGreen line (Aghabi *et al*., 2023). After transfection, parasites were cultured in D3 media containing 5 mM FUDR for 5 days to select for disruption of the *uprt* locus as described previously (Shen *et al*., 2014). Following selection, tdTomato parasites were collected by FACS using an Aria III (BD Biosciences), sorted directly into a 96 well plate pre-seeded with HFFs and incubated at 37°C with 5% CO2 for 5-7 days. Positive fluorescent plaques were then screened by PCR for the disruption of the *uprt* locus and the correct insertion of the reporter cassette (primers oHL247 and oHL248). Fluorescent parasites were resorted as above to ensure purity of populations. For all sgRNA sequences and primers see **Table S10.**

### Chemicals

*N,N,N’,N’*-tetrakis-(2-pyridylmethyl)ethylenediamine (TPEN) (Merck, 616394), made up in DMSO and used at 5 µM. Tetrathiomolybdate (TTM), made up in PBS and used at 25 µM (Merck, 323446). 2,2’-bipyridine (BIP) (VWR, SIALD216305), used at 100 µM, pyridoxal isonicotinoyl hydrazone (PIH) (abcam, ab145871) used at 100 µM, Deferoxamine mesylate salt (Sigma, D9533), concentration as indicated in the text.

### Transcriptomics analysis using bulk RNA-sequencing

T75 flasks of HFF were incubated in either standard D3 or D3 supplemented 100 μM DFO for 24 h prior to infection. Each T75 was then infected with 6-7 × 10^6^ cells of RHΔKu80 parasites and cultured for 24 hours prior to parasite collection. Parasites were pelted by centrifugation at 1500 × g for 10 min, washed and pellets stored at -80 °C until required. RNA was extracted from the pellets using the RNAeasy kit (Qiagen) according to the manufacturer’s instructions. RNA libraries were then prepared using Illumina Stranded mRNA library preparation method and sequenced at 2 × 75 bp to an average of more than 5 million reads per sample. Raw sequencing data (FASTQ format) was processed using the Galaxy public server hosted by EuPathDB (https://veupathdb.globusgenomics.org/). FastQC and Trimmomatic were used for quality control and to remove low quality reads (where Q < 20 across 4 bp sliding windows) and adapter sequences (Bolger, Lohse and Usadel, 2014). The filtered reads were aligned to the *T. gondii* ME49 genome using HISAT2 (Kim, Langmead and Salzberg, 2015). These sequence alignments were used to identify reads uniquely mapped to annotated genes using Htseq-count. Differential expression analysis was performed in R using DESeq2 (Love, Huber and Anders, 2014). Raw FASTQ files will be available from the EBI ENA online server under project PRJEB67890.

### Identification of iron response element-like sequences in the *T. gondii* transcriptome

The *Toxoplasma gondii* reference ME49 transcriptome (v51, ToxoDB.org) was used to identify IRE sequences (Amos *et al*., 2022). At the time of analysis many *T. gondii* transcripts lack annotated UTRs, for these genes a 1kb region up- and down-stream of the transcription start site (Markus *et al*., 2021)/stop codons was included in the bioinformatic search. A 1 kb region was selected based on the average annotated UTR size which we found to be approximately 750 bp as previously reported (Waldman *et al*., 2020). These sequences were then entered into the SIREs online web tool (http://ccbg.imppc.org/sires/), accessed Jan 2023 (Campillos *et al*., 2010). Only sequences with a ‘high’ or ‘medium’ score were selected for further investigation.

### Flow cytometry

Reporter parasites were grown in untreated HFF cells or HFF cells pretreated for 24 h with 500 nM DFO (unless indicated) for 24 h (unless indicated). Cells were washed, parasites mechanically released from host cells and filtered through a 3 mm filter to remove host cell debris. Parasites were pelleted by centrifugation and resuspended in Ringers solution (115 mM NaCl, 3 mM KCl, 2 mM CaCl_2_, 1 mM MgCl_2_, 3 mM NaH_2_PO_4_, 10 mM HEPES, 10 mM glucose), then analysed on a BD Celesta analyser using FACSDiva software (BD Biosciences). Data acquired from at least 30,000 parasites was gated on forward and side scatter and on green fluorescence. All data were analysed using FlowJo v10 (BD Biosciences) and the geometric mean reported.

### Reporter gene expression by microscopy

Confluent HFF cells grown on coverslips were untreated or pretreated for 24 h with 100 mM DFO prior to infection with reporter cell lines. Infected cells were incubated for 24 h before fixation with 4% paraformaldehyde at room temperature for 20 mins. Cells were blocked and permabilised in blocking buffer (2% bovine serum albumin, 0.05% Triton X-100 in PBS) for 20 mins and mounted onto slides using Fluoromount with DAPI (Southern Biotech). Micrograph images were obtained using a DeltaVision widefield (Applied Precision) or Leica DiM8 (Leica Microsystems) microscope and processed using SoftWoRx. Vacuolar tdTomato fluorescence was quantified using a custom ImageJ macro and presented as a ratio compared to mNeonGreen expression.

### Reporter gene expression by RTqPCR

RT-qPCR was used to assay relative *fit* and *tdTomato* expression. HFF cells were untreated or treated with DFO (100 mM) for 24 hours prior to infection with the reporter lines (RHΔKu80, ΔUPRT:tdTomato*_dhfr_*, ΔUPRT:tdTomato*_fit_* or ΔUPRT:tdTomato*_fit_*_Δ*IRE*_). Parasites were cultured as indicated before mechanical lysis of host cells and filtration through a 3 mm to remove host cell debris. Parasites were then pelleted by centrifugation at 1500 × g for 10 min, the media removed and cell pellets stored at -80 °C. Total RNA was extracted from parasite pellets using the RNAeasy Mini kit (Qiagen) and DNAse I 1U/µL (Invitrogen) treated for 15 mins at room temperature followed by DNAse denaturation at 65 °C for 20 mins. cDNA synthesis was performed with High-Capacity cDNA Reverse Transcription Kit (Applied Biosystems) according to manufacturer’s instructions using 1 mg of DNAseI treated RNA. RT-qPCR was carried out on the Applied Biosystems 7500 Real Time PCR system using Power SYBRgreen PCR master mix (Invitrogen), 2 ng of 1:10 diluted cDNAs per reaction and the following cycling conditions: 95 °C for 10 minutes, 40 cycles of 95 °C for 15 seconds, 60 °C for 1 minute. RT-PCR primers sequences can be found in **Table S10.** Reactions were run in triplicate from three independent biological experiments. Relative fold changes for treated vs. untreated cells was calculated using the Pfaffl method (Pfaffl, 2001) using *actin* as a housekeeping gene control. Graphpad Prism 9 was used to perform statistical analysis.

### Fluorescent parasite growth assay

Iron chelators (DFO, PIH and BIP) were tested for parasite inhibition as previously described (Aghabi *et al*., 2023) (Aghabi *et al*., 2023). Briefly, confluent HFF monolayers grown in clear bottomed 96 well plates were infected with 5000 ΔKu80:tdTomato (Aghabi *et al*., 2023) parasites/well Invasion was allowed to occur for 2 h before the media was removed, and indicated chelator concentrations added and serially diluted. Cells were incubated for 4 days before parasite proliferation quantified by PheraStar plate reader. Results were normalised to untreated wells and inhibition curves plotted by non-linear regression using Prism (v10).

### Aconitase activity assay

Aconitase activity was quantified based the production of NADPH from *T. gondii* (Quirós, 2018b). Untreated parasites or those treated for 24 h with DFO were harvested as described above, pelleted by centrifugation and resuspended in aconitase activity buffer (50 mM Tris–HCl, pH 7.4, 6 mM sodium citrate, 0.2% Triton X-100, 0.6 mM MnCl_2_) before lysis by three rounds of freeze-thaw. Protein content of lysates was quantified by Bradford assay (ThermoScientific, B6916) following manufacturer’s instructions. 10 mg of parasite lysate was combined with 0.2 mM NADP and 0.004 U of isocitrate dehydrogenase (Sigma, <u>I2002</u>) in a total volume of 100 ml . In each assay, 10 µl of disrupted and filtered host cell debris was included to ensure no contamination by host cell enzymes present. The absorbance was read at 340 nm at 25 °C every 5 mins for 1 h on a PheraStar plate reader. After normalization to blanks, the slope was calculated from the linear section of the curve and protein activity (c) calculated using c = slope_final_/ε · b with ε of NADP = 6.22 mM^−1^ cm^−1^ and path length (b) set to 1. Results represent four independent experiments performed in quadruplicate.

### Immunoprecipitation of *T. gondii* proteins and interacting RNAs

Parasites (RHDku80:mNeonGreen-Ty or RH,ACN-Ty) were cultured for 18 hours in HFFs which had been pre-treated in D3 with 100 mM DFO for 24 hours. To collect the parasites host cell monolayers were scraped, passed through a syringe with 23G gauge needle three times, filtered through a 3 mM polycarbonate filter membranes to remove host debris and then pelleted by centrifugation. Cell pellets were frozen and stored at -80°C prior to further processing. For immunoprecipitation, 5 × 10^8^ parasites were lysed for 30 minutes on ice in 500 mL lysis buffer (50 mM Tris pH7.4, 150 mM NaCl, 1 mM EDTA, 1 mM EGTA, 1 % Triton-X100 with Halt protease inhibitor (ThermoFisher, 78429) and SUPERase RNAse inhibitors 1 U/mL). Lysates were centrifuged at 13,000 rpm at 4°C for 15 mins and the supernatants (cleared lysate) blocked for 1 h at 4°C with 50 mL of Pierce Protein-G magnetic beads with end-over-end rotation. Lysates were then incubated for 3 h at 4 °C with 50 mL of Pierce Protein-G beads bound with mouse anti-Ty antibody (Invitrogen Ty1 Tag Monoclonal BB2). Beads were washed 6 times with wash buffer (200 mM Tris pH 9, 100 mM Potassium acetate, 0.5 % (v/v) Triton x-100, 1 mM EGTA, 0.1 % Tween 20, 2.5 mM Dithiothreitol). Total RNA was then extracted from the washed beads by Trizol-chloroform extraction, following manufacturer’s instructions. Extracted RNA was treated with DNAse I (Invitrogen) as above before proceeding to cDNA synthesis using High-Capacity cDNA Reverse Transcription Kit (Applied Biosystems), according to the manufacturer’s instructions. Resulting cDNAs were diluted 1:5 prior to qPCR which was performed as described above. For primer sequences see **Table S10.**

### mRNA stability assay

HFF cells were pre-treated for 24 hours with either fresh D3 or D3 with 100 mM DFO before being infected with tdTomato*_dhfr_*or tdTomato*_fit_* reporter parasites. At 18 h post infection, 10mg/ml Actinomycin D was added to the culture media at in the treatment dishes. Parasites were collected at 0, 1, 3 and 5 hours post Actinomycin D treatment. At each collection, host cells were mechanically lysed and filtered through a 3mm membrane to remove debris. Parasites were pelleted by centrifugation and immediately frozen on dry ice for storage at -80°C. RNA was extracted using a Qiagen RNAeasy kit according to manufacturers instructions. RNAs were the processed as described above for RTqPCR. All primer sequences can be found in **Table S10.** All data was plotted in Prism (v10) and non-linear regression (one phase decay) performed to determine mRNA half-life.

## Supporting information

Tabls S1

## Supplementary Figure Legends

**Figure S1 – PCR confirmation of iron reporter lines. A.** Schematic of the cloning strategy for adding the reporter cassettes into the *uprt* locus using CRISPR-Cas9 **B-D.** PCRs showing successful amplification of the full reporter tdTomato*_dhfr_* (**B**), tdTomato*_fit_* (**C**), tdTomato*_fit_*_Δ_*_IRE_* (**C**), and tdTomato_iscu_ (**D**), cassettes into the *uprt* locus of RHΔKu80:mNeonGreen parasites. Parental line included as negative control **E.** Overlapping histogram showing tdTomato fluorescence in untreated tdTomato*_dhfr_*(black), tdTomato*_iscu_* (blue) and tdTomato*_fit_* (red) reporter lines as measured by flow cytometry.

**Figure S2 – IRP1 is highly conserved between apicomplexans and mammals. A.** Amino acid sequence alignment of the *Toxoplasma gondii* aconitase hydratase ACN/IRP (TGME49_226730), *Plasmodium falciparum* aconitase hydratase (PF3D7_1342100) and human IRP1 (UniProt: P21399) and IRP2 (UniProt: P48200) in ClustalW format made using T-Coffee (Notredame, Higgins and Heringa, 2000). Green triangles indicate the residues interacting with the FeS cluster. Purple triangles indicate residues shown to interact with ferritin mRNAs in human IRP1 (Walden *et al*., 2006). **B.** Alphafold (Jumper *et al*., 2021; Varadi *et al*., 2022) structure prediction showing high structural conservation between TgACN (magenta) and HsIRP1(pdb 2B3X) (cyan), including around the FeS coordination core (yellow, inset).

**Figure S3 – Acontiase associates with *fit* transcripts. A.** Normalised counts for *fit* and *rng2* transcripts from RNAseq dataset comparing parasites after 24 hours treatment with 100 µM DFO, compared to untreated parasites. Points represent 3 independent experiments, bars at meanLJ±LJSD. Two (**B** and **C**) further biological replicates showing CAN-Ty interacts with *fit* mRNA. Blots showing the immunoprecipitation of mNeonGreen-Ty and aconitase-Ty from *T. gondii* parasites. Wash – the output of the 6^th^ and final washing step. CDPK1 included as cytosolic control. From the corresponding pull down, DNA-agarose gel showing qPCR products amplified from reverse transcribed RNA from either lysates or RNA co-precipitated with mNeonGreen-Ty and ACN-Ty proteins.

## Acknowledgements

We would like to acknowledge Aarti Krishnan and Dominique Soldati for the iACN:Ty line and Lilach Sheiner for the anti-TOM40 antibody. Additionally, we would like to thank Shikha Shikha for assistance with AlphaFold and Kasia Moderynska for her guidance and RIP-qPCR protocol. M.A.S is funded by an Early Career Award from the Wellcome Trust (225677/Z/22/Z). C.R.H. is funded by a Sir Henry Dale Fellowship from the Wellcome Trust and the Royal Society (213455/Z/18/Z).

## Supplementary Tables

**Table S1** – RNAseq results from experiment comparing parasites cultured for 24 hours in 100µM DFO to untreated parasites.

**Table S2** – Genes predicted by MetalPredator to encode proteins with iron sulphur clusters. Log_2_ fold changes and adjusted *p* values from RNAseq experiment comparing parasites cultured for 24 hours in 100µM DFO to untreated parasites.

**Table S3** - FeS synthesis pathway genes (Pamukcu *et al*., 2021b) with log_2_ fold changes and adjusted *p* values from RNAseq data.

**Table S4** - Genes upregulated in *T. gondii* bradyzoites (log_2_ fold changes > 2, adjusted *p* value < 0.05) from Waldman *et al*. 2020 in low iron conditions

**Table S5** – Raw output file from SIREs web server (Campillos *et al*., 2010) identifying putative IRE-like sequences in curated *T. gondii* transcriptome.

**Table S6** – Curated list of genes containing putative IRE-like sequences in the *T. gondii* transcriptome containing only high quality and medium quality scoring sequences as identified using SIREs (Campillos *et al*., 2010).

**Table S7** - Curated list of genes containing putative IRE-like sequences in the *T. gondii* transcriptome containing only high quality and medium quality scoring sequences, limited to those found within untranslated regions (Campillos *et al*., 2010).

**Table S8** - Curated list of genes containing putative IRE-like sequences in the *T. gondii* transcriptome containing only high quality and medium quality scoring sequences with from motif classes I, II and VIII, found within the untranslated regions - as identified using SIREs (Campillos *et al*., 2010).

**Table S9** – RTqPCR Ct values from RIP-qPCR experiments to look for transcripts associated with *Toxoplasma gondii* aconitase.

**Table S10** - Oligonucleotides used in this study.

## References

Aghabi, D. et al. (2023) ‘The vacuolar iron transporter mediates iron detoxification in Toxoplasma gondii’, Nature Communications, 14(1), p. 3659. Available at: 10.1038/s41467-023-39436-y.

Alén, C. and Sonenshein, A.L. (1999) ‘Bacillus subtilis aconitase is an RNA-binding protein’, Proceedings of the National Academy of Sciences, 96(18), pp. 10412–10417. Available at: 10.1073/pnas.96.18.10412.

Alvarez, H.M. et al. (2010) ‘Tetrathiomolybdate Inhibits Copper Trafficking Proteins Through Metal Cluster Formation’, Science, 327(5963), pp. 331–334. Available at: 10.1126/science.1179907.

Amos, B. et al. (2022) ‘VEuPathDB: the eukaryotic pathogen, vector and host bioinformatics resource center’, Nucleic Acids Research, 50(D1), pp. D898–D911. Available at: 10.1093/nar/gkab929.

Andrews, S.C., Robinson, A.K. and Rodríguez-Quiñones, F. (2003) ‘Bacterial iron homeostasis’, FEMS Microbiology Reviews, 27(2–3), pp. 215–237. Available at: 10.1016/S0168-6445(03)00055-X.

Arnaud, N. et al. (2007) ‘The iron-responsive element (IRE)/iron-regulatory protein 1 (IRP1)- cytosolic aconitase iron-regulatory switch does not operate in plants’, The Biochemical Journal, 405(3), pp. 523–531. Available at: 10.1042/BJ20061874.

Barylyuk, K. et al. (2020) ‘A Comprehensive Subcellular Atlas of the Toxoplasma Proteome via hyperLOPIT Provides Spatial Context for Protein Functions’, Cell Host & Microbe, 28(5), pp. 752–766.e9. Available at: 10.1016/j.chom.2020.09.011.

Bergmann, A. et al. (2020) ‘Toxoplasma gondii requires its plant-like heme biosynthesis pathway for infection’, PLOS Pathogens, 16(5), p. e1008499. Available at: 10.1371/journal.ppat.1008499.

Bohne, W. and Roos, D.S. (1997) ‘Stage-specific expression of a selectable marker in Toxoplasma gondii permits selective inhibition of either tachyzoites or bradyzoites’, Molecular and Biochemical Parasitology, 88(1), pp. 115–126. Available at: 10.1016/S0166-6851(97)00087-X.

Bolger, A.M., Lohse, M. and Usadel, B. (2014) ‘Trimmomatic: a flexible trimmer for Illumina sequence data’, Bioinformatics, 30(15), pp. 2114–2120. Available at: 10.1093/bioinformatics/btu170.

Calla-Choque, J.S. et al. (2014) ‘α-Actinin TvACTN3 of Trichomonas vaginalis is an RNA-Binding Protein That Could Participate in Its Posttranscriptional Iron Regulatory Mechanism’, BioMed Research International, 2014, p. e424767. Available at: 10.1155/2014/424767.

Campillos, M. et al. (2010) ‘SIREs: searching for iron-responsive elements’, Nucleic Acids Research, 38(suppl_2), pp. W360–W367. Available at: 10.1093/nar/gkq371.

Carbajo, C.G. et al. (2021) ‘Novel aspects of iron homeostasis in pathogenic bloodstream form Trypanosoma brucei’, PLOS Pathogens, 17(6), p. e1009696. Available at: 10.1371/journal.ppat.1009696.

Casey, J.L. et al. (1988) ‘Two genetic loci participate in the regulation by iron of the gene for the human transferrin receptor’, Proceedings of the National Academy of Sciences of the United States of America, 85(6), pp. 1787–1791. Available at: 10.1073/pnas.85.6.1787.

DeRusso, P.A. et al. (1995) ‘Expression of a Constitutive Mutant of Iron Regulatory Protein 1 Abolishes Iron Homeostasis in Mammalian Cells (∗)’, Journal of Biological Chemistry, 270(26), pp. 15451–15454. Available at: 10.1074/jbc.270.26.15451.

Donald, R.G. and Roos, D.S. (1995) ‘Insertional mutagenesis and marker rescue in a protozoan parasite: cloning of the uracil phosphoribosyltransferase locus from Toxoplasma gondii.’, Proceedings of the National Academy of Sciences, 92(12), pp. 5749–5753. Available at: 10.1073/pnas.92.12.5749.

Erlitzki, R., Long, J.C. and Theil, E.C. (2002) ‘Multiple, Conserved Iron-responsive Elements in the 31-Untranslated Region of Transferrin Receptor mRNA Enhance Binding of Iron Regulatory Protein 2*’, Journal of Biological Chemistry, 277(45), pp. 42579–42587. Available at: 10.1074/jbc.M207918200.

Fontenot, C.R. and Ding, H. (2023) ‘Ferric uptake regulator (Fur) binds a [2Fe-2S] cluster to regulate intracellular iron homeostasis in Escherichia coli’, Journal of Biological Chemistry, 299(6), p. 104748. Available at: 10.1016/j.jbc.2023.104748.

Fox, B.A., Gigley, J.P. and Bzik, D.J. (2004) ‘Toxoplasma gondii lacks the enzymes required for de novo arginine biosynthesis and arginine starvation triggers cyst formation’, International Journal for Parasitology, 34(3), pp. 323–331. Available at: 10.1016/j.ijpara.2003.12.001.

Gao, F. and Dubos, C. (2021) ‘Transcriptional integration of plant responses to iron availability’, Journal of Experimental Botany, 72(6), pp. 2056–2070. Available at: 10.1093/jxb/eraa556.

Garber, I. and Pudek, M. (2014) ‘A novel deletion in the iron-response element of the L-ferritin gene, causing hyperferritinaemia cataract syndrome’, Annals of Clinical Biochemistry, 51(6), pp. 710–713. Available at: 10.1177/0004563214542289.

Garza, K.R. et al. (2020) ‘Differential translational control of 5’ IRE-containing mRNA in response to dietary iron deficiency and acute iron overload’, Metallomicsr?: integrated biometal science, 12(12), pp. 2186–2198. Available at: 10.1039/d0mt00192a.

Goncalves, S. et al. (2008) ‘Deferiprone targets aconitase: implication for Friedreich’s ataxia treatment’, BMC neurology, 8, p. 20. Available at: 10.1186/1471-2377-8-20.

Gunawardena, N.D. et al. (2016) ‘Aconitase: An Iron Sensing Regulator of Mitochondrial Oxidative Metabolism and Erythropoiesis’, Blood, 128(22), p. 74. Available at: 10.1182/blood.V128.22.74.74.

Harding, C.R. et al. (2020) ‘Genetic screens reveal a central role for heme metabolism in artemisinin susceptibility’, Nature Communications, 11(1), p. 4813. Available at: 10.1038/s41467-020-18624-0.

Heaslip, A.T. et al. (2011) ‘The Motility of a Human Parasite, Toxoplasma gondii, Is Regulated by a Novel Lysine Methyltransferase’, PLOS Pathogens, 7(9), p. e1002201. Available at: 10.1371/journal.ppat.1002201.

Hentze, M.W. et al. (1987) ‘A cis-acting element is necessary and sufficient for translational regulation of human ferritin expression in response to iron’, Proceedings of the National Academy of Sciences of the United States of America, 84(19), pp. 6730–6734. Available at: 10.1073/pnas.84.19.6730.

Ihara, F. and Nishikawa, Y. (2014) ‘Starvation of low-density lipoprotein-derived cholesterol induces bradyzoite conversion in Toxoplasma gondii’, Parasites & Vectors, 7, p. 248. Available at: 10.1186/1756-3305-7-248.

Jumper, J. et al. (2021) ‘Highly accurate protein structure prediction with AlphaFold’, Nature, 596(7873), pp. 583–589. Available at: 10.1038/s41586-021-03819-2.

Kato, J. et al. (2007) ‘Iron/IRP-1-dependent regulation of mRNA expression for transferrin receptor, DMT1 and ferritin during human erythroid differentiation’, Experimental Hematology, 35(6), pp. 879–887. Available at: 10.1016/j.exphem.2007.03.005.

Katris, N.J. et al. (2014) ‘The Apical Complex Provides a Regulated Gateway for Secretion of Invasion Factors in Toxoplasma’, PLOS Pathogens, 10(4), p. e1004074. Available at: 10.1371/journal.ppat.1004074.

Kim, D., Langmead, B. and Salzberg, S.L. (2015) ‘HISAT: a fast spliced aligner with low memory requirements’, Nature Methods, 12(4), pp. 357–360. Available at: 10.1038/nmeth.3317.

Koeller, D. M. et al. (1989) ‘A cytosolic protein binds to structural elements within the iron regulatory region of the transferrin receptor mRNA’, Proceedings of the National Academy of Sciences of the United States of America, 86(10), pp. 3574–3578. Available at: 10.1073/pnas.86.10.3574.

Koeller, D M et al. (1989) ‘A cytosolic protein binds to structural elements within the iron regulatory region of the transferrin receptor mRNA.’, Proceedings of the National Academy of Sciences, 86(10), pp. 3574–3578. Available at: 10.1073/pnas.86.10.3574.

León-Sicairos, C.R. et al. (2023) ‘The Non-Canonical Iron-Responsive Element of IRE-tvcp12 Hairpin Structure at the 31-UTR of Trichomonas vaginalis TvCP12 mRNA That Binds TvHSP70 and TvACTN-3 Can Regulate mRNA Stability and Amount of Protein’, Pathogens, 12(4), p. 586. Available at: 10.3390/pathogens12040586.

Li, L. et al. (2008) ‘Yap5 Is an Iron-Responsive Transcriptional Activator That Regulates Vacuolar Iron Storage in Yeast’, Molecular and Cellular Biology, 28(4), pp. 1326–1337. Available at: 10.1128/MCB.01219-07.

Li, Yaqiong et al. (2023) ‘Rapid metabolic reprogramming mediated by the AMP-activated protein kinase during the lytic cycle of Toxoplasma gondii’, Nature Communications, 14(1), p. 422. Available at: 10.1038/s41467-023-36084-0.

Licon, M.H. et al. (2023) ‘A positive feedback loop controls Toxoplasma chronic differentiation’, Nature Microbiology, 8(5), pp. 889–904. Available at: 10.1038/s41564-023-01358-2.

Lind, M.I. et al. (2006) ‘Of two cytosolic aconitases expressed in Drosophila, only one functions as an iron-regulatory protein’, The Journal of Biological Chemistry, 281(27), pp. 18707–18714. Available at: 10.1074/jbc.M603354200.

Love, M.I., Huber, W. and Anders, S. (2014) ‘Moderated estimation of fold change and dispersion for RNA-seq data with DESeq2’, Genome Biology, 15(12), p. 550. Available at: 10.1186/s13059-014-0550-8.

Loyevsky, M. et al. (2001) ‘An IRP-like protein from Plasmodium falciparum binds to a mammalian iron-responsive element’, Blood, 98(8), pp. 2555–2562. Available at: 10.1182/blood.V98.8.2555.

Loyevsky, M. et al. (2003) ‘Expression of a recombinant IRP-like Plasmodium falciparum protein that specifically binds putative plasmodial IREs’, Molecular and Biochemical Parasitology, 126(2), pp. 231–238. Available at: 10.1016/S0166-6851(02)00278-5.

MacRae, J.I. et al. (2012) ‘Mitochondrial metabolism of glucose and glutamine is required for intracellular growth of Toxoplasma gondii’, Cell Host & Microbe, 12(5), pp. 682–692. Available at: 10.1016/j.chom.2012.09.013.

Markus, B.M. et al. (2021) ‘High-Resolution Mapping of Transcription Initiation in the Asexual Stages of Toxoplasma gondii’, Frontiers in Cellular and Infection Microbiology, 10. Available at: https://www.frontiersin.org/articles/10.3389/fcimb.2020.617998 (Accessed: 4 August 2023).

Marondedze, C. et al. (2016) ‘The RNA-binding protein repertoire of Arabidopsis thaliana’, Scientific Reports, 6(1), p. 29766. Available at: 10.1038/srep29766.

Martínez-Pastor, M.T. and Puig, S. (2020) ‘Adaptation to iron deficiency in human pathogenic fungi’, Biochimica et Biophysica Acta (BBA) - Molecular Cell Research, 1867(10), p. 118797. Available at: 10.1016/j.bbamcr.2020.118797.

Morel, J.-D. et al. (2022) ‘The mouse metallomic landscape of aging and metabolism’, Nature Communications, 13(1), p. 607. Available at: 10.1038/s41467-022-28060-x.

Notredame, C., Higgins, D.G. and Heringa, J. (2000) ‘T-coffee: a novel method for fast and accurate multiple sequence alignment11Edited by J. Thornton’, Journal of Molecular Biology, 302(1), pp. 205–217. Available at: 10.1006/jmbi.2000.4042.

Oexle, H., Gnaiger, E. and Weiss, G. (1999) ‘Iron-dependent changes in cellular energy metabolism: influence on citric acid cycle and oxidative phosphorylation’, Biochimica et Biophysica Acta (BBA) - Bioenergetics, 1413(3), pp. 99–107. Available at: 10.1016/S0005-2728(99)00088-2.

Owen, D. and Kühn, L.C. (1987) ‘Noncoding 3’ sequences of the transferrin receptor gene are required for mRNA regulation by iron’, The EMBO journal, 6(5), pp. 1287–1293. Available at: 10.1002/j.1460-2075.1987.tb02366.x.

Pamukcu, S. et al. (2021a) ‘Differential contribution of two organelles of endosymbiotic origin to iron-sulfur cluster synthesis and overall fitness in Toxoplasma’, PLoS pathogens, 17(11), p. e1010096. Available at: 10.1371/journal.ppat.1010096.

Pamukcu, S. et al. (2021b) ‘Differential contribution of two organelles of endosymbiotic origin to iron-sulfur cluster synthesis and overall fitness in Toxoplasma’, PLOS Pathogens, 17(11), p. e1010096. Available at: 10.1371/journal.ppat.1010096.

Pfaffl, M.W. (2001) ‘A new mathematical model for relative quantification in real-time RT– PCR’, Nucleic Acids Research, 29(9), p. e45.

Philpott, C.C., Klausner, R.D. and Rouault, T.A. (1994) ‘The bifunctional iron-responsive element binding protein/cytosolic aconitase: the role of active-site residues in ligand binding and regulation.’, Proceedings of the National Academy of Sciences of the United States of America, 91(15), pp. 7321–7325.

Pino, P. et al. (2007) ‘Dual Targeting of Antioxidant and Metabolic Enzymes to the Mitochondrion and the Apicoplast of Toxoplasma gondii’, PLOS Pathogens, 3(8), p. e115. Available at: 10.1371/journal.ppat.0030115.

Plata-Guzmán, L.Y. et al. (2023) ‘Stem-Loop Structures in Iron-Regulated mRNAs of Giardia duodenalis’, International Journal of Environmental Research and Public Health, 20(4), p. 3556. Available at: 10.3390/ijerph20043556.

Quirós, P.M. (2018a) ‘Determination of Aconitase Activity: A Substrate of the Mitochondrial Lon Protease’, in S. Cal and A.J. Obaya (eds) Proteases and Cancer: Methods and Protocols. New York, NY: Springer (Methods in Molecular Biology), pp. 49–56. Available at: 10.1007/978-1-4939-7595-2_5.

Quirós, P.M. (2018b) ‘Determination of Aconitase Activity: A Substrate of the Mitochondrial Lon Protease’, in S. Cal and A.J. Obaya (eds) Proteases and Cancer: Methods and Protocols. New York, NY: Springer (Methods in Molecular Biology), pp. 49–56. Available at: 10.1007/978-1-4939-7595-2_5.

Reinert, A. et al. (2019) ‘Iron concentrations in neurons and glial cells with estimates on ferritin concentrations’, BMC Neuroscience, 20(1), p. 25. Available at: 10.1186/s12868-019-0507-7.

Sahu, T. et al. (2014) ‘ZIPCO, a putative metal ion transporter, is crucial for Plasmodium liver-stage development’, EMBO Molecular Medicine, 6(11), pp. 1387–1397. Available at: 10.15252/emmm.201403868.

Sanchez, M. et al. (2007) ‘Iron-regulatory proteins limit hypoxia-inducible factor-2alpha expression in iron deficiency’, Nature Structural & Molecular Biology, 14(5), pp. 420–426. Available at: 10.1038/nsmb1222.

Sanchez, M. et al. (2011) ‘Iron regulatory protein-1 and -2: transcriptome-wide definition of binding mRNAs and shaping of the cellular proteome by iron regulatory proteins’, Blood, 118(22), pp. e168–e179. Available at: 10.1182/blood-2011-04-343541.

Schaefer-Ramadan, S. et al. (2019) ‘Synthesis of TPEN variants to improve cancer cells selective killing capacity’, Bioorganic Chemistry, 87, pp. 366–372. Available at: 10.1016/j.bioorg.2019.03.045.

Shen, B. et al. (2014) ‘Efficient Gene Disruption in Diverse Strains of Toxoplasma gondii Using CRISPR/CAS9’, mBio, 5(3), p. 10.1128/mbio.01114-14. Available at: 10.1128/mbio.01114-14.

Shin, L.-J. et al. (2013) ‘IRT1 DEGRADATION FACTOR1, a RING E3 Ubiquitin Ligase, Regulates the Degradation of IRON-REGULATED TRANSPORTER1 in Arabidopsis’, The Plant Cell, 25(8), pp. 3039–3051. Available at: 10.1105/tpc.113.115212.

Sidik, S.M. et al. (2016) ‘A Genome-wide CRISPR Screen in Toxoplasma Identifies Essential Apicomplexan Genes’, Cell, 166(6), pp. 1423–1435.e12. Available at: 10.1016/j.cell.2016.08.019.

Sloan, M.A., Aghabi, D. and Harding, C.R. (2021) ‘Orchestrating a heist: uptake and storage of metals by apicomplexan parasites’, Microbiology, 167(12), p. 001114. Available at: 10.1099/mic.0.001114.

Smith, T.A. et al. (2022) ‘Screening the Toxoplasma kinome with high-throughput tagging identifies a regulator of invasion and egress’, Nature Microbiology, 7(6), pp. 868–881. Available at: 10.1038/s41564-022-01104-0.

Tang, Y. and Guest, J.R. (1999) ‘Direct evidence for mRNA binding and post-transcriptional regulation by Escherichia coli aconitases’, Microbiology, 145(11), pp. 3069–3079. Available at: 10.1099/00221287-145-11-3069.

Valasatava, Y. et al. (2016) ‘MetalPredator: a web server to predict iron–sulfur cluster binding proteomes’, Bioinformatics, 32(18), pp. 2850–2852. Available at: 10.1093/bioinformatics/btw238.

Varadi, M. et al. (2022) ‘AlphaFold Protein Structure Database: massively expanding the structural coverage of protein-sequence space with high-accuracy models’, Nucleic Acids Research, 50(D1), pp. D439–D444. Available at: 10.1093/nar/gkab1061.

Volz, K. (2021) ‘Conservation in the Iron Responsive Element Family’, Genes, 12(9), p. 1365. Available at: 10.3390/genes12091365.

Walden, W.E. et al. (2006) ‘Structure of Dual Function Iron Regulatory Protein 1 Complexed with Ferritin IRE-RNA’, Science, 314(5807), pp. 1903–1908. Available at: 10.1126/science.1133116.

Waldman, B.S. et al. (2020) ‘Identification of a Master Regulator of Differentiation in Toxoplasma’, Cell, 180(2), pp. 359–372.e16. Available at: 10.1016/j.cell.2019.12.013.

Wang, J. and Pantopoulos, K. (2011) ‘Regulation of cellular iron metabolism’, Biochemical Journal, 434(3), pp. 365–381. Available at: 10.1042/BJ20101825.

White, M.W. and Suvorova, E.S. (2018) ‘Apicomplexa Cell Cycles: Something Old, Borrowed, Lost, and New’, Trends in Parasitology, 34(9), pp. 759–771. Available at: 10.1016/j.pt.2018.07.006.

Zhu, Y. et al. (2023) ‘Iron stress affects the survival of Toxoplasma gondii’. Research Square. Available at: 10.21203/rs.3.rs-3240882/v1.

