## Supplemental figures for "Keeping FIT: Iron-mediated post-transcriptional regulation in *Toxoplasma gondii*"

### Figure S1

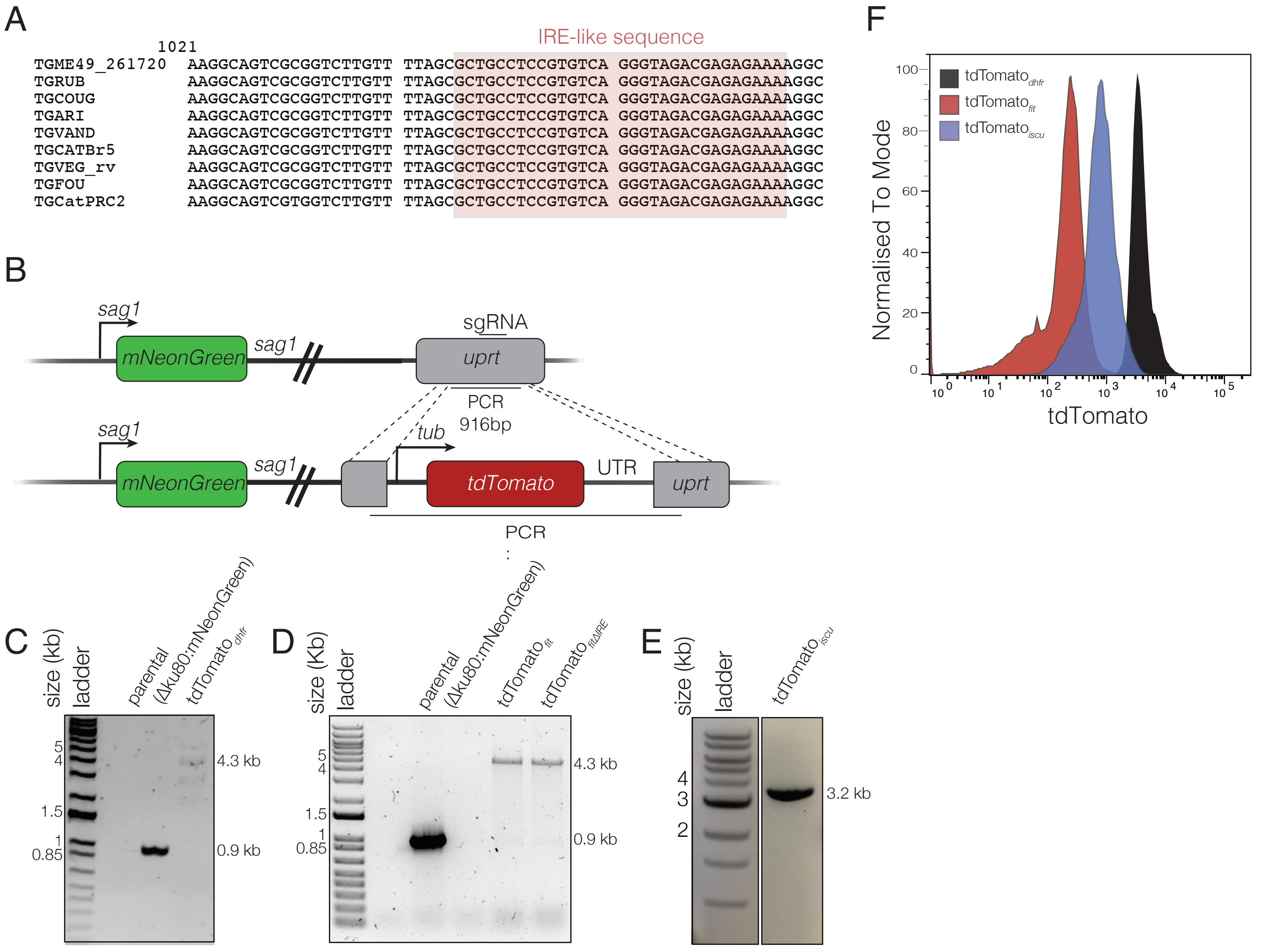


**Figure S1 – PCR confirmation of iron reporter lines. A.** Alignment of *fit* UTR from selected *T. gondii* strains. **B.** Schematic of the cloning strategy for adding the reporter cassettes into the *uprt* locus using CRISPR-Cas9 **C-E**. PCRs showing successful amplification of the full reporter $tdTomato_{dhfr}$ (**C**), $tdTomato_{fit}$ (**D**), $tdTomato_{fit\Delta IRE}$ (**D**), and $tdTomato_{iscu}$ (**E**), cassettes into the *uprt* locus of RHΔKu80:mNeonGreen parasites. Parental line included as negative control **F**. Overlapping histogram showing tdTomato fluorescence in untreated $tdTomato_{dhfr}$ (black), $tdTomato_{iscu}$ (blue) and $tdTomato_{fit}$ (red) reporter lines as measured by flow cytometry.

### Figure S2

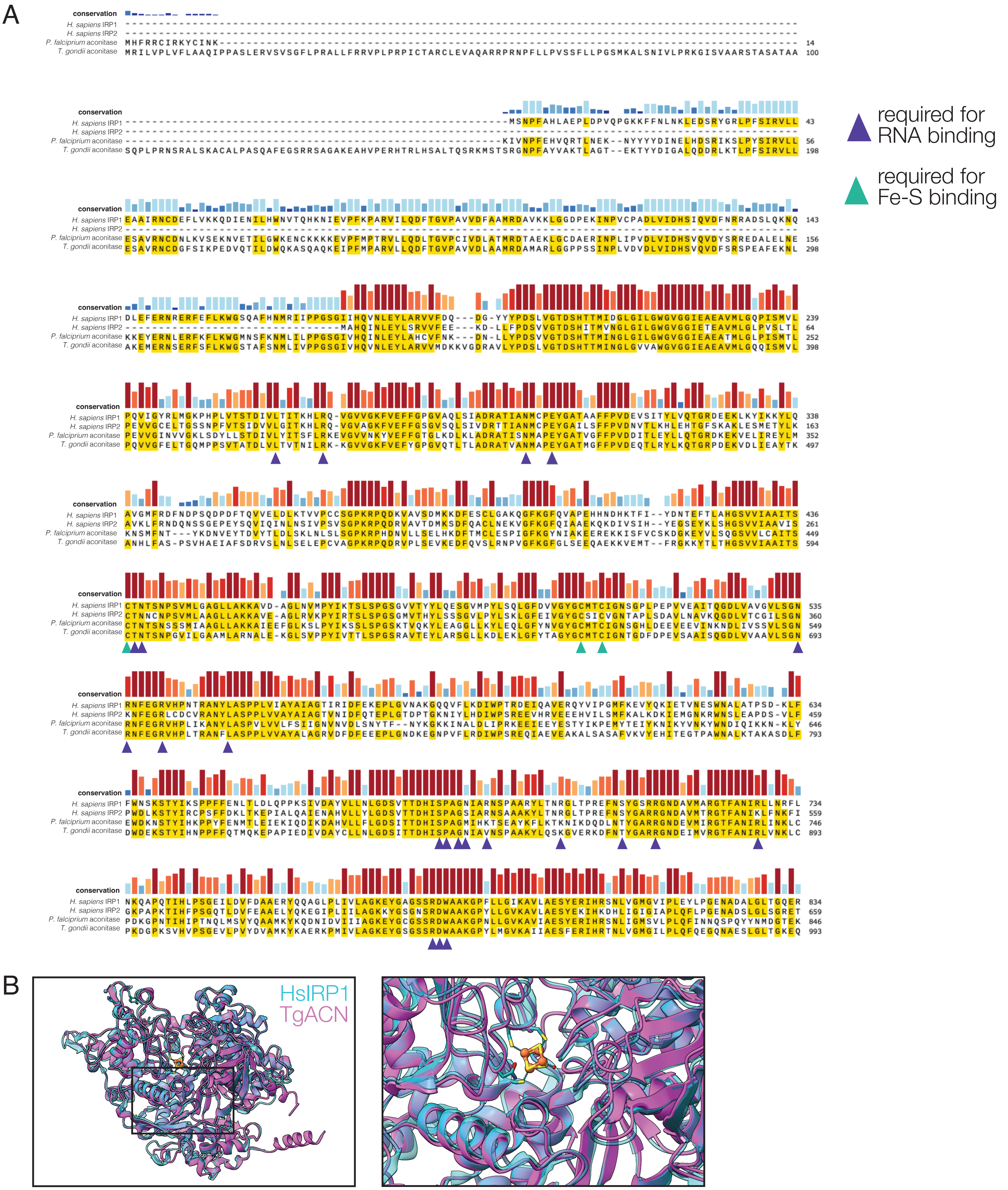


**Figure S2 – IRP1 is highly conserved between apicomplexans and mammals. A.** Amino acid sequence alignment of the *Toxoplasma gondii* aconitase hydratase ACN/IRP (TGME49_226730), *Plasmodium falciparum* aconitase hydratase (PF3D7_1342100) and human IRP1 (UniProt: P21399) and IRP2 (UniProt: P48200) in ClustalW format made using T-Coffee (Notredame, Higgins and Heringa, 2000). Green triangles indicate the residues interacting with the FeS cluster. Purple triangles indicate residues shown to interact with ferritin mRNAs in human IRP1 (Walden et al., 2006). **B.** Alphafold (Jumper et al., 2021; Varadi et al., 2022) structure prediction showing high structural conservation between TgACN (magenta) and HsIRP1(pdb 2B3X) (cyan), including around the FeS coordination core (yellow, inset).

Figure S3

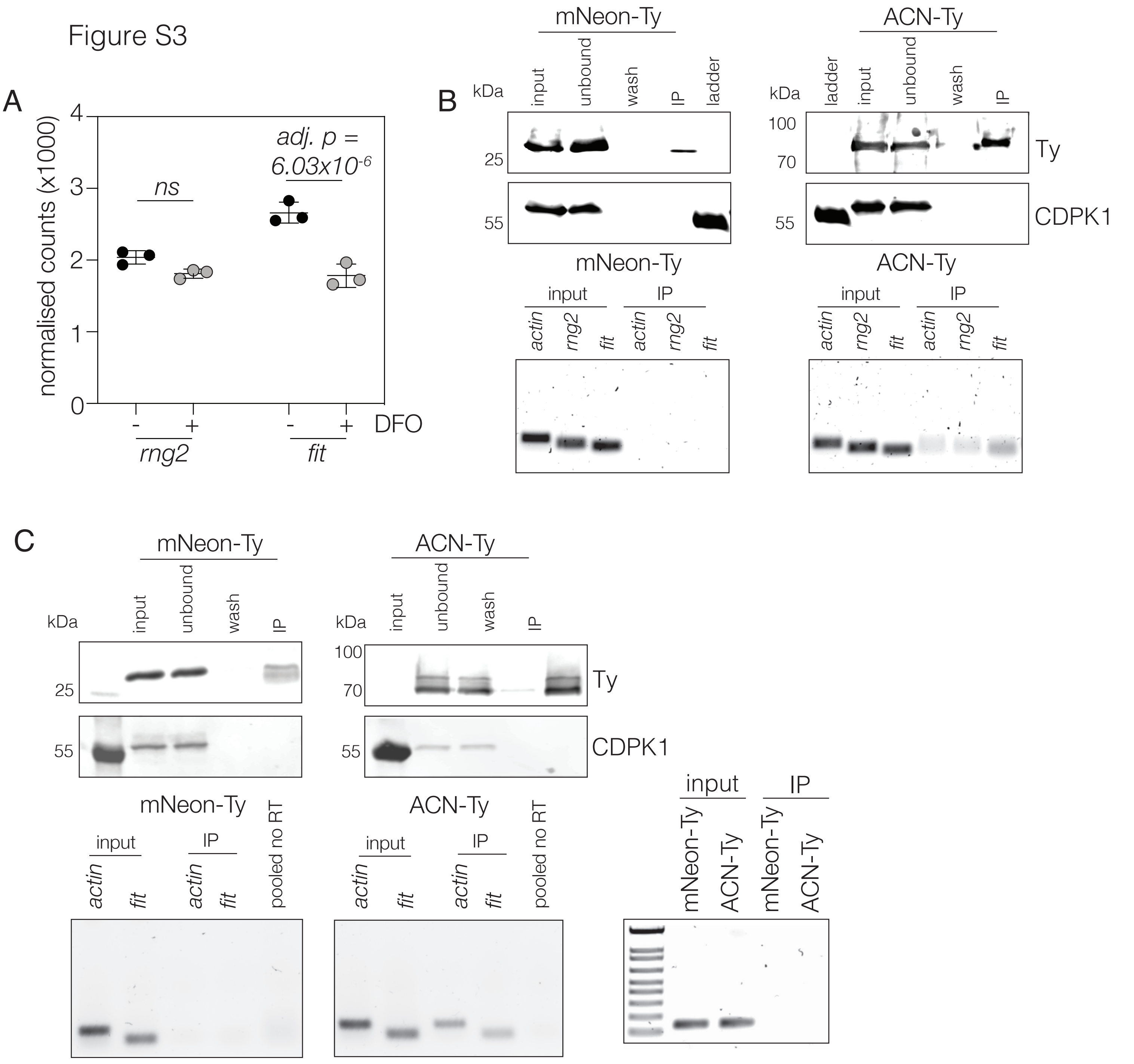


**Figure S3 – Acontiase associates with fit transcripts. A.** Normalised counts for *fit* and *rng2* transcripts from RNAseq dataset comparing parasites after 24 hours treatment with 100 µM DFO, compared to untreated parasites. Points represent 3 independent experiments, bars at mean ± SD. Two (**B and C**) further biological replicates showing CAN-Ty interacts with *fit* mRNA. Blots showing the immunoprecipitation of mNeonGreen-Ty and aconitase-Ty from *T. gondii* parasites. Wash – the output of the 6th and final washing step. CDPK1 included as cytosolic control. From the corresponding pull down, DNA-agarose gel showing qPCR products amplified from reverse transcribed RNA from either lysates or RNA co-precipitated with mNeonGreen-Ty and ACN-Ty proteins.
